# A novel machine learning-based algorithm for eQTL identification reveals complex pleiotropic effects in the MHC region

**DOI:** 10.1101/2025.05.06.652558

**Authors:** Ronnie Y. Li, Chang Su, Zhaohui S. Qin

## Abstract

Expression quantitative trait loci (eQTLs) are regulatory variants that affect the expression level of their target genes and have significant impact on disease biology. However, eQTL mapping has been done mostly in one tissue at a time, despite the known prevalence of correlations among tissues. Multivariate analyses incorporating multiple phenotypes are available, but they emphasize linear combinations of phenotypes. We present MTClass, a machine learning framework that attempts to classify an individual’s genotype based on a vector of multi-phenotype expression levels of a given gene. We conduct simulation studies and multiple case studies using real and imputed data, and we demonstrate that MTClass detects more functionally relevant variants and genes compared to existing single-tissue approaches as well as multi-phenotype association tests. Our results suggest that the importance of expression regulation at the MHC region may have been underestimated, and they provide fresh biological insights into genetic variants that have pleiotropic effects, influencing gene expression in a complex manner.

**Key points:**

1. MTClass is a machine learning-based approach that classifies genotypes based on multi-phenotype expression data, providing a novel method for identifying eQTLs.
2. MTClass outperforms traditional linear methods like MultiPhen and MANOVA in detecting eQTLs with greater functional impact and in capturing complex genotype-phenotype relationships.
3. MTClass identified immune-related variants in the HLA region, suggesting that existing approaches may have underestimated the complexity of these variants’ effects across tissues.
4. MTClass is more flexible and reliable than linear multivariate methods, handling multicollinearity, zero-expressed features, and various input values with greater ease.

## Introduction

Evaluating the consequences of genetic variation on the biological functioning on cells, tissues, and disease risks is an important undertaking in genetics. Widely adopted genome-wide association studies (GWASs) have revealed that most variants associated with a given phenotype are in noncoding regions of the genome^1,2^, leading to the hypothesis that these variants affect regulatory functions instead of protein-coding ones. For example, Kapoor and colleagues identified 210 common risk variants for QT interval variation; strikingly, all of them were noncoding^3^. Similar evidence has been found for other phenotypes like restless leg syndrome^4^ and schizophrenia^5^. Indeed, such discoveries have spawned a new field in human genetics studying expression quantitative trait loci (eQTLs)^6^.

An eQTL is a genetic variant that partially explains the variance in the expression level of a gene either proximally (in *cis*) or distally (in *trans*). Standard eQTL mapping involves a direct association test between the genotypes of a variant and a gene’s expression levels. Mapping and linkage studies have consistently shown that eQTLs have considerable biological significance^6^. GWAS-significant single nucleotide polymorphisms (SNPs) are more likely to colocalize with eQTLs^7^. Additionally, eQTLs are more likely to have regulatory implications such as altering transcription factor binding affinity^8^. They can also be organized into tissue-specific modular networks to inform disease risk and coherent biological processes^9^.

Despite the widespread significance of eQTLs, gene expression levels are tissue-specific, and most existing eQTL studies have focused on identifying eQTLs one tissue at a time (*i.e.*, single-tissue approach). However, somatic tissues are statistically and biologically correlated^10^, raising the possibility that an eQTL can affect gene expression in more than one tissue. Nevertheless, current approaches to detect the pleiotropic interaction between one genotype and multiple phenotypes are relatively limited, and most of them run a “reverse” regression. Notable examples include MultiPhen^11^, MANOVA^12^, SCOPA/META-SCOPA^13^, MASH^14^, and others^15–18^. Originally designed for GWAS, MultiPhen performs ordinal regression on the phenotypes, testing the linear combination of phenotypes that best explains the given genotype. Similarly, SCOPA and META-SCOPA both run a reverse linear regression. One significant limitation facing these methods is that they only consider linear effects of genotype on phenotype, which is rather restrictive.

In this study, we present MTClass, a model-free approach adopting an ensemble machine learning framework to classify a genotype based on multivariate gene expression levels from multiple sources (**Figure 1**). Instead of calculating a p-value under the traditional statistical testing framework, we rank eGene-eQTL pairs based on classification performance. In doing so, we hypothesize that variants with superior classification performance will be more likely to be multi-phenotype eQTLs, as they exhibit a detectable mathematical relationship between genotype and phenotypes. We enumerate cases to which we apply MTClass on real and simulated data and compare our method to state-of-the-art existing approaches, demonstrating that our top identified variants are likely to have more functional relevance. Further, we extend our algorithm to classify genotypes based on expression levels from multiple exons, as we believe that exon-level expression vector can provide richer detail into gene expression than an aggregated single measure of expression. Next, we use a multi-layer perceptron (MLP) to classify genotypes based on the two-dimensional matrices of exon-level and tissue-level expression of a gene. Lastly, we utilize MTClass on two additional datasets with greater sample sizes, one from the PsychENCODE Consortium^19–21^ and a single-cell RNA-seq dataset from the Human Cell Atlas^22^. We compare our approach to state-of-the-art existing methods, and we illustrate interesting anecdotal examples of multi-phenotype eQTLs.

**Figure 1.**
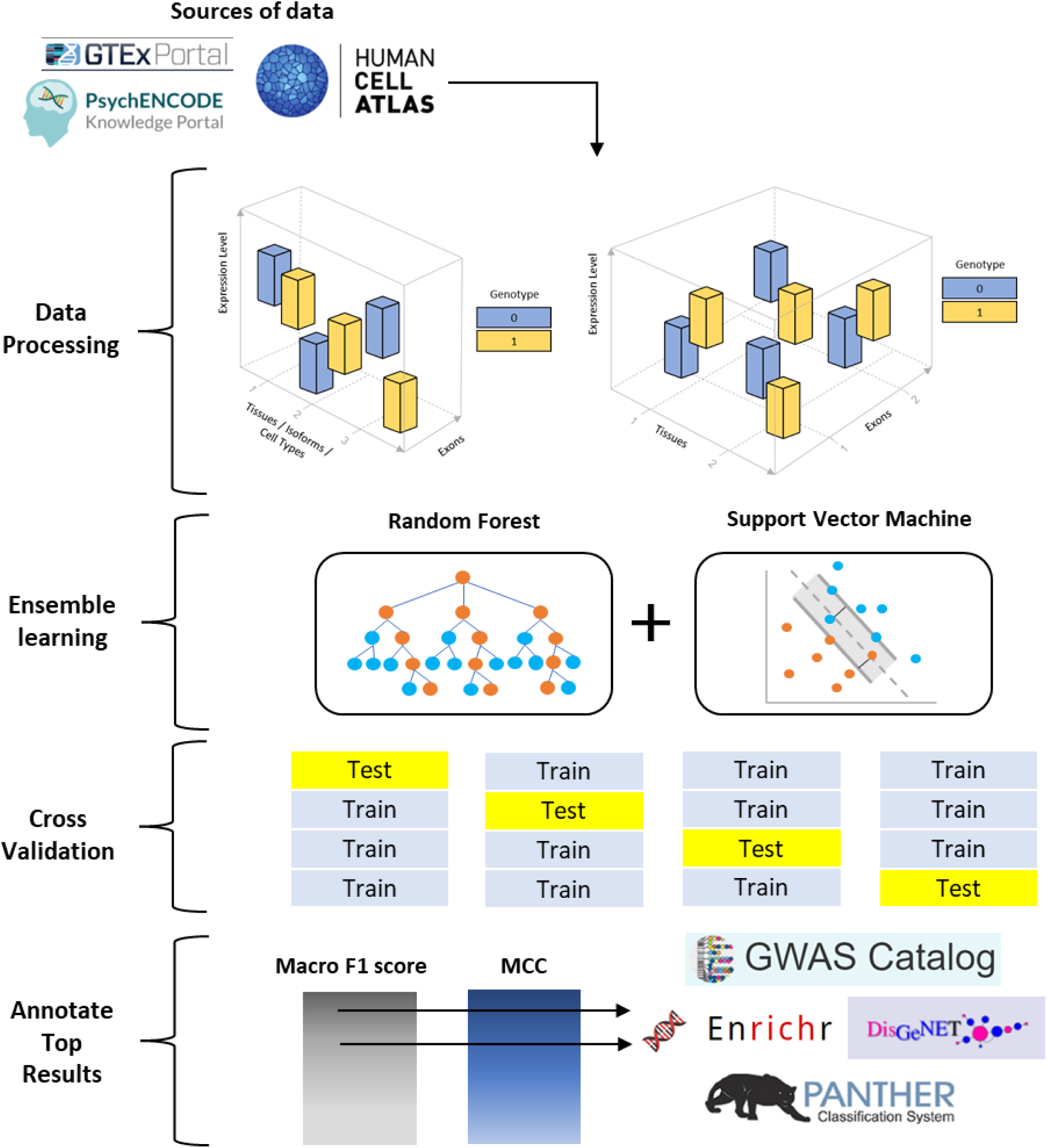
Schematic overview of MTClass algorithm. After processing the data from the GTEx Consortium, the PsychENCODE Consortium, and the Human Cell Atlas, MTClass uses an ensemble learning framework comprised of Random Forest and Support Vector Machine, combined with 4-fold cross validation, to classify genotypes according to the vector of expression levels from a nearby gene. Variants are then sorted by macro F1 or MCC, and the top variants are annotated using various bioinformatic databases.

## Results

### Simulation study

To demonstrate the utility of MTClass in detecting nonlinear relationships between genotypes and phenotypes, we present two simulated datasets. For both datasets, we assume a sample size of 200: 100 dominant genotypes (denoted as 0) and 100 heterozygous/recessive genotypes (denoted as 1). In the first dataset, we simulated 3 tissue-level expression measures (“dimensions”) in spherical patterns, such that samples with the 0 genotype have expression levels on the surface of a sphere of radius 5, while samples with the 1 genotype have expression levels on the surface of a sphere of radius 20. Effectively, the two genotypes have expression levels with the same mean but different variances **(Supplementary Figure 1A-1C)**. This makes the separation boundary between the two genotypes nonlinear. In the second dataset, the three-dimensional expression levels followed interlocking sinusoidal patterns **(Supplementary Figure 1D-1F)**. We tested MultiPhen and MANOVA on both datasets and found that neither method was able to output a significant p-value on either dataset. For the first and second datasets, MultiPhen reported p-values of 0.540 and 0.997, while MANOVA reported p-values of 0.460 and 0.997, respectively. However, our ensemble approach was able to classify the genotypes well, achieving macro F1 scores of 0.95 and 0.925, and MCC scores of 0.905 and 0.851, respectively. **Supplementary Figure 1** depicts the area under the receiver operating characteristic (AUROC) curves of our ensemble classifier, showing that the ensemble approach can robustly detect such nonlinear relationships whereas existing methods cannot.

### GTEx data

A significant obstacle in mapping eQTLs in multiple tissues is limited tissue availability. The GTEx Consortium has exerted remarkable effort in overcoming this challenge by collecting many tissues from the same donors and performing rigorous quality control on the specimens^23^. We used publicly available normalized multi-tissue gene expression data in transcripts per million (TPM) obtained from the GTEx Consortium, version 8, consisting of a total of 948 donors across 54 tissues, although the exact number of donors per tissue varied considerably. We also utilized the publicly available exon-level expression data across 13 brain tissues. The genotype data of for the GTEx donors were accessed through dbGaP^24^ and converted into PLINK^25^ file formats for downstream analyses. **Table 1** lists the sample sizes, number of genes, variants, and features used for each study.

**Table 1.** Number of unique eGenes, variants, donors, and features used in all experiments. Features refer to tissues (multi-tissue study), exons (multi-exon study), both tissues and exons (2D study), isoforms (isoQTL study), or cell types (OneK1K study).

| Experiment | Genes | Variants | Sample size | Number of features |
| --- | --- | --- | --- | --- |
| Multi-tissue studies |  |  |  |  |
| 9-tissue | 13811 | 3013552 | 103 | 9 |
| Brain tissue | 13812 | 3017214 | 317 | 13 |
| 48-tissue | 13813 | 3018097 | 838 | 48 |
| Multi-exon studies |  |  |  |  |
| Brain_Amygdala | 5040 | 1854146 | 129 | Varies by gene |
| Brain_Anterior_cingulate_cortex_BA24 | 5045 | 1855259 | 147 | Varies by gene |
| Brain_Caudate_basal_ganglia | 5049 | 1859142 | 194 | Varies by gene |
| Brain_Cerebellar_Hemisphere | 5044 | 1842548 | 175 | Varies by gene |
| Brain_Cerebellum | 5051 | 1861331 | 209 | Varies by gene |
| Brain_Cortex | 5050 | 1853926 | 205 | Varies by gene |
| Brain_Frontal_Cortex_BA9 | 5050 | 1860706 | 175 | Varies by gene |
| Brain_Hippocampus | 5049 | 1860066 | 165 | Varies by gene |
| Brain_Hypothalamus | 5050 | 1852746 | 170 | Varies by gene |
| Brain_Nucleus_accumbens_basal_ganglia | 5052 | 1849364 | 202 | Varies by gene |
| Brain_Putamen_basal_ganglia | 5049 | 1859289 | 170 | Varies by gene |
| Brain_Spinal_cord_cervical_c-1 | 5041 | 1846537 | 126 | Varies by gene |
| Brain_Substantia_nigra | 5037 | 1848812 | 114 | Varies by gene |
| Other studies |  |  |  |  |
| 2D (multi-tissue/multi-exon) | 10227 | 3857513 | 103 | Varies by gene |
| isoQTL (PsychENCODE) | 5435 | 1265533 | 900 | Varies by gene |
| OneK1K (scRNA-seq) | 9237 | 1251974 | 982 | 7 |

### Multi-tissue study

We enumerate three cases with different subsets of donors and tissues to demonstrate the utility of our approach: the 9-tissue case, the brain tissue case, and the 48-tissue case. The 9-tissue case is carefully selected such that there is no missing data, while the brain tissue case and 48-tissue case contained a substantial amount of missing data.

### 9-tissue case

The 9-tissue case consisted of gene-level expression data from 9 tissue types and across 103 donors. The nine tissues included subcutaneous adipose, tibial artery, lung, skeletal muscle, tibial nerve, skin (not sun-exposed, suprapubic), skin (sun-exposed, lower leg), thyroid, and whole blood. We chose these 9 tissues to maximize the number of tissues while maintaining sufficient sample sizes without missing data. To ensure that our results were reproducible and consistent, we ran our ensemble classifier three times on the same dataset, each time with different random seeds, then aggregated the macro F1 scores and MCC by taking the median values across the three iterations. **Figure 2A** illustrates the Manhattan plot for the 9-tissue case, using the median macro F1 score as the classification performance metric. Switching to median MCC to rank the top variants and genes triggered minimal change to the results, as the two metrics are highly correlated **(Supplementary Figure 2)**. A total of 54 eGene-eQTL pairs belonging to 8 genes achieved both perfect median F1 scores and median MCC in the three iterations of MTClass (**Table 2**). For the purposes of comparison, we also ran MultiPhen and MANOVA on the same datasets. We noticed that there was a notable degree of overlap among the top 1000 and top 5000 variants selected by all three methods. Among the three top 1000 variants lists, 373 variants (37.3%) were common to all three methods, and among the top 5000 variant lists, there were 2913 common variants (58.3%, **Supplementary Figure 3**). The top-performing eQTLs of the top 10 eGenes ranked by MCC from the 9-tissue case are summarized in **Table 3**.

**Figure 2.**
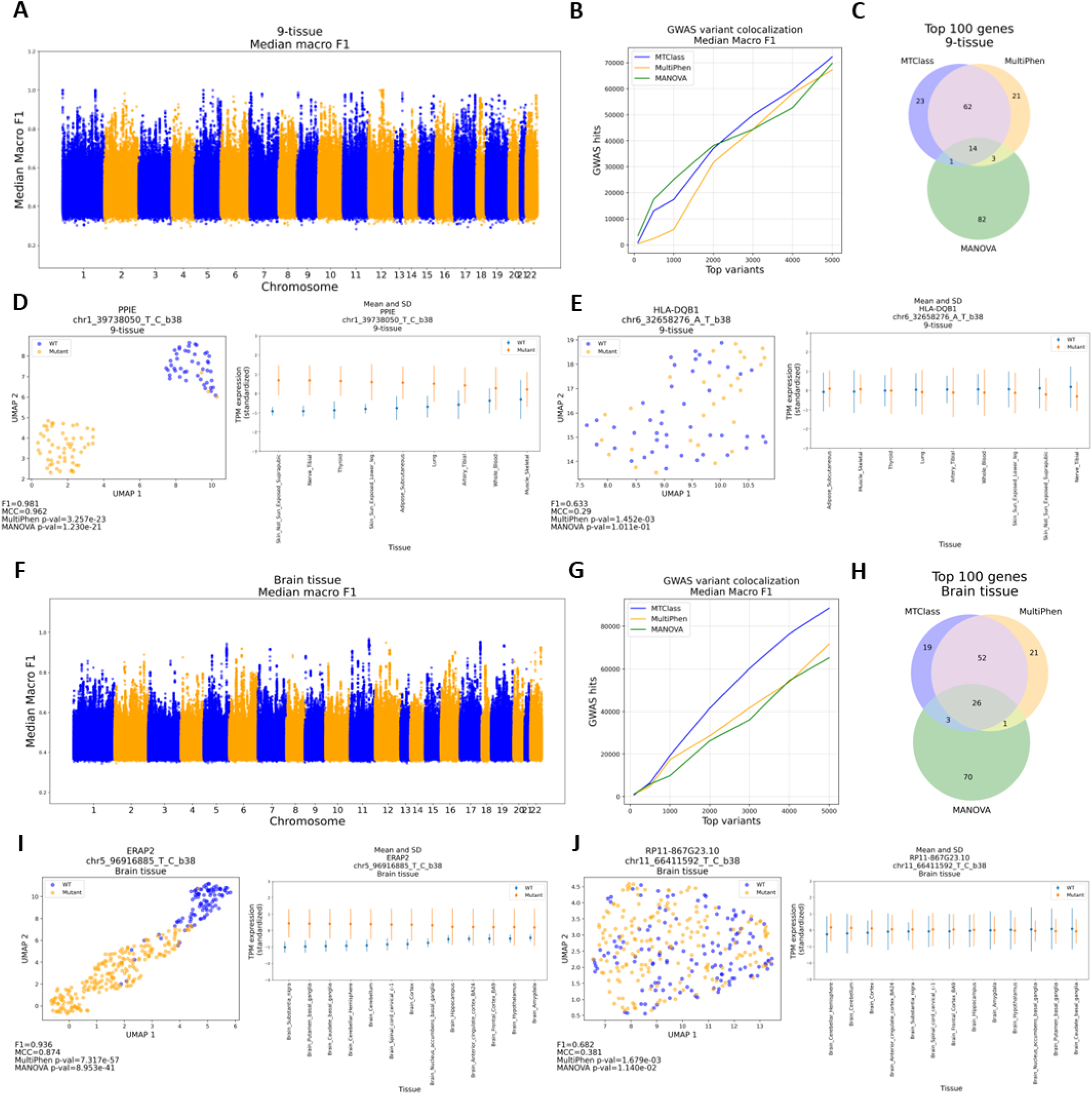
Functional assessment of top variants identified by MTClass compared to MultiPhen and MANOVA in the multi-tissue study. **A-E** refer to the 9-tissue case, and **F-J** refer to the brain tissue case. **A**. Manhattan plot of variants according to their median macro F1 score. **B.** GWAS variant colocalization analysis of top variants from each method. **C.** Overlap of top 100 genes detected by each method. **D.** PPIE-chr1_39738050_T_C_b38 (rs12074147) was reported as highly significant by all three methods. UMAP analysis of this pair (left) and the expression levels across 9 tissues (right). **E.** HLA-DQB1-chr6_32658276_A_T_b38 (rs28633132) was classified well by MTClass but not reported as significant by MultiPhen and MANOVA. **F.** Manhattan plots of variants by median macro F1 score. **G.** GWAS variant colocalization analysis of top variants. **H.** Overlap of top 100 genes detected by each method. **I.** ERAP2-chr5_96916885_T_C_b38 (rs2910686) was reported as highly significant by MultiPhen and MANOVA. UMAP analysis of this pair (left) and the expression levels across 13 brain tissues (right). **J.** RP11-867G23.10-chr11_66411592_T_C_b38 (rs12785651) was classified well by MTClass but not reported as significant by MultiPhen and MANOVA.

**Table 2.** The 54 eGene-eQTL pairs that achieved perfect classification performance (median macro F1 score and median MCC) in three iterations of MTClass in the 9-tissue case. These eGenes are enriched for housekeeping processes involving DNA metabolism and processing, as well as cell division processes.

| **BTNL3** | **CLECL12A** | **DDX11** | **HORMAD1** |
| --- | --- | --- | --- |
| rs72494581 | rs111930678, rs113362278<br>rs12230638, rs12231872<br>rs199560361, rs200911123<br>rs201541579, rs61913537 | chr12_31080603_GA_G_b38<br>rs10843879, rs11051236<br>rs2543261, rs2553134<br>rs2553147, rs3881296<br>rs3930906, rs4031323<br>rs4031341, rs4031360<br>rs7133363, rs71440936<br>rs7309520 | chr1_150730558_CAGA_C_b38<br>rs112718432, rs11803940<br>rs11807450, rs1336899<br>rs1336900, rs2184833<br>rs3768005, rs6700022<br>rs868751 |
| **PDK1P5** | **PPIE** | **RPS26** | **ZNF300P1** |
| rs4781810<br>rs62043986 | chr1_39739455_TAAAG_T_b38<br>rs1046988, rs11809515<br>rs12094291, rs2274949<br>rs3738673, rs41267029<br>rs56064244, rs58589165<br>rs6664570, rs6670841<br>rs7513045, rs7520588<br>rs7547787, rs7548120<br>rs9787249 | rs1131017 | rs12659118, rs76433514 |

**Table 3.**
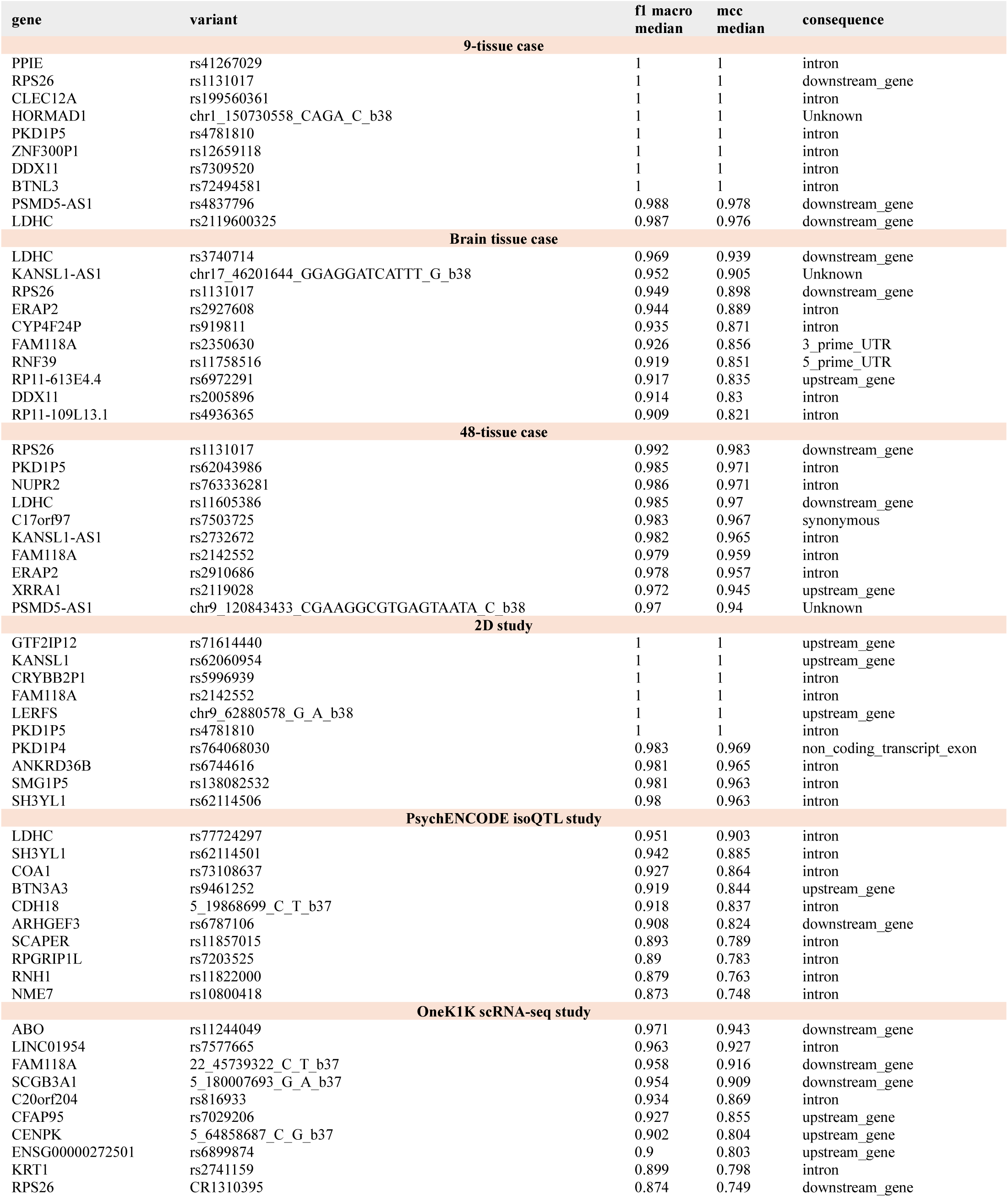
The top-performing eQTLs from the top 10 eGenes by MCC in the 9-tissue case, the brain tissue case, and the 48-tissue case (multi-tissue study), the PsychENCODE isoQTL study, and the OneK1K scRNA-seq study.

Next, we tested whether the top variants identified by MTClass were more functionally relevant compared to the top variants identified by MultiPhen and MANOVA. To do this, we downloaded the entire GWAS Catalog^26^ and examined whether each method’s top variants were colocalized with known trait-associated SNPs from the catalog. We therefore defined “GWAS hits” as the total number of unique trait-associated SNPs within 10 kilobases of a given variant’s genomic position. After selecting the same number of top variants from MultiPhen and MANOVA by p-value, we calculated the GWAS hits for the top 5000 variants selected by each method.

We demonstrated that the top variants selected by MTClass colocalized with more known GWAS-identified trait-associated SNPs than did the top variants selected by MultiPhen, regardless of whether we sorted by macro F1 or MCC. This superior performance was preserved when we aggregated the macro F1 score across 3 iterations by calculating the median (**Figure 2B**). MTClass performed similarly to MANOVA overall in terms of GWAS hits. However, MTClass outperformed MANOVA in the latter half of the variants tested. These results suggest that the MTClass-identified variants have more potential functional relevance than those identified by MultiPhen.

We also determined whether the genes detected by our method were more functionally impactful. To this end, we selected the top 100 genes from each method. We ran a Gene Ontology (GO)^27^ statistical overrepresentation test on the genes on PantherDB^28^, using the 13,810 total genes as the reference list. We restricted the GO analysis to GO terms with at most 50 genes in the reference list to avoid detecting overly generic GO terms. In accordance with the null hypothesis, we found no significant differences in nominal p-values among the three groups as measured by a one-way ANOVA. Among the top 100 genes, only 14 genes (14%) were common to all three methods, suggesting that each method largely detected distinct patterns in the data (**Figure 2C**).

### Examples from the 9-tissue case

Altogether, there were a total of 54 variants belonging to 8 genes that achieved a perfect median macro F1 score and median MCC across all three iterations of MTClass (**Table 2**). We analyzed the functional potential for these 8 genes using PantherDB^28^ and Enrichr^29^, and we discovered that they were enriched for DNA metabolism and processing, as well as cell division processes **(Supplementary Figure 4)**.

We delved more deeply into the eGene-eQTL pairs that were classified very well by MTClass. One of the eQTL-eGene pairs with a high macro F1 score and MCC across all 3 iterations of the 9-tissue case was PPIE-chr1_39738050_T_C_b38 (rs12074147), which achieved a median macro F1 score of 0.981 and a median MCC of 0.962. This pair was also deemed highly significant by MultiPhen (*p* = 3.26 × 10^−23^) and MANOVA (*p* = 1.23 × 10^−21^). Indeed, dimensionality reduction using Uniform Manifold Approximation and Projection (UMAP) shows high separability between the two genotypes (**Figure 2D**). According to the GTEx Consortium, this SNP is a significant eQTL in 49 out of 54 tissues (90.7%), including all 9 of the tissues in this study.

Moreover, the PPIE gene is expressed highly in many tissues, especially testis, EBV-transformed lymphocytes, and the tibial nerve. PPIE is a cyclophilin, which means it participates in many important biological processes, including mitochondrial metabolism, apoptosis, and inflammation^30^. Diseases associated with PPIE include ischemic reperfusion injury, influenza, and certain cancers^31^. Biologically, given the plethora of functions served by PPIE, it is reasonable that this eQTL would be significant in many somatic tissues.

Next, we surveyed the eQTLs that were classified well by MTClass (median F1 > 0.60) but not by MultiPhen or MANOVA (*p* > 0.001). An interesting example of such an eQTL is HLA-DQB1-chr6_32658276_A_T_b38 (rs28633132). For this eGene-eQTL pair, MTClass reported a median macro F1 score of 0.633 and median MCC of 0.290, while MultiPhen reported a p-value of 1.45 × 10^−3^, and MANOVA reported a p-value of 1.01 × 10^−1^. UMAP dimensionality reduction analysis shows that the two genotype groups are not easily separable, suggesting a more complex relationship that would be better captured by MTClass (**Figure 2E**). This gene plays a critical role in the immune system, as it is the human version of MHC class II. This SNP is a significant eQTL in 19 out of 54 GTEx tissues (35.2%), including 5 out of the 9 tissues (55.6%) that were tested in this case. Thus, it is conceivable that this eGene-eQTL pair would have far-reaching effects as a multi-tissue eQTL.

### Brain tissue case

There are 13 brain tissues profiled in the GTEx Consortium: amygdala, anterior cingulate cortex (Brodmann area 24), caudate nucleus, cerebellar hemisphere, cerebellum, cortex, frontal cortex (Brodmann area 9), hippocampus, hypothalamus, nucleus accumbens, putamen, spinal cord (cervical C1), and substantia nigra. The v8 version of the GTEx data contains 317 donors contributing at least one tissue type. The average percentage of missing data across all 13 tissues was 47% (between 33-64%, **Supplementary Figure 5)**.

Expression levels from missing donor-tissue combinations were imputed for each gene using predictive mean matching with chained equations^32^. We implemented the imputation strategy five times and took the arithmetic mean (**Supplementary Figure 6**).

To improve reproducibility and mitigate uncertainty in our results, we ran our ensemble classifier three times, using different initialization weights each time, and we computed the median macro F1 score and median MCC for each eGene-eQTL pair. Due to large amounts of missing data, the overall classification performance in the brain tissue case was poorer compared to that in the 9-tissue case, as evidenced by the Manhattan plots (**Figure 2F**). Although there were no eGene-eQTL pairs in the brain tissue case that achieved a median macro F1 or MCC equal to 1.0, we still observed evidence of strong classification performance among the top variants.

As in the 9-tissue case, we tested whether the top variants and genes identified by MTClass were more functionally relevant compared to those identified by other methods. We computed the colocalization with known GWAS trait-associated SNPs within a 10 kb neighborhood for each top variant as we did in the 9-tissue case, and we found that the top variant from MTClass had substantially more colocalization with known trait-associated SNPs compared to those from MultiPhen and MANOVA (**Figure 2G**). The top-performing eQTLs of the top 10 eGenes ranked by MCC from the brain tissue case are summarized in **Table 3**. Among the top 100 eGenes detected by MTClass, MultiPhen, and MANOVA, 26 genes were common to all three methods (**Figure 2H**).

### Examples from the brain tissue case

One eGene-eQTL pair that was identified in the brain tissue case was ERAP2-chr5_96916885_T_C_b38 (rs2910686), which achieved a median macro F1 of 0.936 and median MCC of 0.874 across 3 iterations. This pair was also reported as highly significant by MultiPhen (*p* = 7.32 × 10^−57^) and MANOVA (*p* = 8.95 × 10^−41^). UMAP dimensionality reduction analysis shows highly separable genotypes (**Figure 2I**). According to the GTEx Consortium, this SNP is a significant eQTL in 49 out of 54 GTEx tissues (90.1%), including all 13 brain tissues tested. Although this gene’s expression profile is not specific to the brain, ERAP2 does play important roles in innate immunity and major histocompatibility complex class I (MHC Class I) antigen processing and presentation^33^. This suggests that ERAP2 might contribute to widespread immune processes that can ultimately impact the brain.

Next, we examined the eGene-eQTL pairs that were classified well by MTClass (median macro F1 score > 0.60) but showed weak associations according to MultiPhen and MANOVA (*p* > 0.001). An interesting eQTL we discovered was RP11-867G23.10-chr11_66411592_T_C_b38 (rs12785651). MTClass reported a median macro F1 score of 0.682 and median MCC of 0.381, while MultiPhen reported a p-value of (1.68 × 10^−3^) and MANOVA reported a p-value of (1.14 × 10^−2^). The genotypes are not separable in UMAP space, but there are subtle differences in expression levels between the two genotypes that MTClass was still able to capture (**Figure 2J**). RP11-867G23.10 is also known as NPAS4, and it is a transcription factor expressed in neurons that regulates the excitatory-inhibitory balance in the brain ^34^. This eQTL is a significant eQTL in 3 out of 13 brain tissues (23.0%) according to the GTEx Consortium: cerebellum, cerebellar hemisphere, and cortex. As a transcriptional regulator, NPAS4 potentially has overarching effects on brain function, but the relationship between genotype and phenotype might be a more nonlinear one, as evidenced by MTClass’ differential ability to detect it.

### 48-tissue case

The 48-tissue case was comprised of expression level vectors from 48 somatic tissues (including all 13 brain tissues), across 838 donors. The goal of this case study was to identify the most broad-acting eQTLs that exert effects across a very large number of tissues. We excluded tissues that had fewer than 100 donors available, thus reducing the total number of tissues from 54 to 48. Across the 48 tissues, the average missing data rate was 62%, ranging from 15% (skeletal muscle) to 85% (uterus and substantia nigra). As with the brain tissue case, we imputed missing TPM measures for each gene using 5 iterations of predictive mean matching using chained equations, then taking the average of the predictions.

We repeated the GWAS variant colocalization analysis in the 48-tissue case and compared our results to MultiPhen and MANOVA. Although MTClass consistently outperformed MultiPhen in terms of neighboring GWAS hits, it only outperformed MANOVA in the latter portion of the top variants tested (top 2500-5000 variants; **Supplementary Figure 7**). We also observed that among the top 100 genes, there was greater overlap among the three methods compared to both the 9-tissue and brain tissue cases, suggesting that the increase in dimensionality tends to cause the methods to converge in their top genes. The top-performing eQTLs of the top 10 eGenes ranked by MCC from the 48-tissue case are summarized in **Table 3**.

Lastly, for all three multi-tissue cases, we utilized the GWAS variant colocalization approach to compare MTClass with the top single-tissue eQTLs from the individual tissues tested. We found that only in the brain tissue case, MTClass outperformed all single-tissue eQTLs in terms of GWAS hits **(Supplementary Figure 8A-8C)**. However, MTClass clearly outperforms random subsets of variants in all three cases **(Supplementary Figure 8D-8F)**. We hypothesize that the reason for MTClass’ relative underperformance in the 9-tissue and 48-tissue cases is the remarkable diversity of the tissues. In the brain tissue case, expression measures from each tissue were highly correlated with the others (*R*^2^ ≥ 0.80), whereas in the other two cases, there was at least one tissue with weak correlations (*R*^2^ ≤ 0.20) to the other tissues. Together, the results from these three studies reinforce the notion that MTClass can be a useful tool to detect multi-tissue eQTLs that have important functional consequences.

### Multi-exon study in brain tissue

Alternative splicing is a major regulator of gene expression, particularly in the human brain^35,36^. Studies have shown that aberrations in alternative splicing contribute to many disorders, including frontotemporal dementia^37^, Duchenne muscular dystrophy^38^, and cystic fibrosis^39^. By studying multi-exon eQTLs, we hope to identify variants that affect the expression of multiple exons within the same gene and thereby contribute to alternative splicing mechanisms or differential expression of transcript isoforms. Identifying these genetic variants can provide insight into both regulation of gene expression and alternative splicing, as exon-level expression might be more informative compared to gene-level expression^40,41^.

To accomplish this goal, we adapted our multi-tissue approach so that the features in our model were expression levels, in TPM, from various exons of the same gene in a single tissue. To maximize the chance of novel discovery, we focused on exon-rich genes defined to be genes containing at least 10 exons. We excluded eGene-eQTL pairs that were not able to be processed by MultiPhen or MANOVA. Altogether, we processed about 5000 genes per tissue (the exact number of genes varied slightly by tissue and is listed in **Table 1**). We ran three iterations of MTClass in each of 13 brain tissues obtained from the GTEx Consortium. We compared our approach to MultiPhen and MANOVA. The top-performing eQTLs of the top 10 eGenes ranked by MCC from each tissue are summarized in **Supplementary Data 1**.

To determine whether the top genes identified by MTClass in each brain tissue were more functionally relevant than those identified by MultiPhen or MANOVA, we focused on four neurological and psychiatric disorders that are known to substantially impact specific areas of the brain: Alzheimer’s Disease (AD) and the amygdala^42,43^, depression and the anterior cingulate cortex^44,45^, schizophrenia and the nucleus accumbens^46,47^, and Parkinson’s Disease (PD) and the substantia nigra^48^. Using version 3 of the DisGeNET database^49^, a comprehensive knowledge base of disease-gene associations, we calculated the amount of overlap between the top genes from each method and known disease-related genes. We selected the top 100 eGenes from the MTClass, MultiPhen, and MANOVA results as well as the top 100 eGenes from the single-tissue eQTL and splicing QTL (sQTL) analyses. Throughout these four comparisons, we found that the top 100 genes detected by MTClass had the greatest overlap with disease-related genes in these three disease-related tissues, outperforming both single-tissue eQTL and sQTL approaches (**Figure 3A**) as well as MultiPhen and MANOVA (**Figure 3B**). We also discovered that the top genes detected by MTClass were largely distinct from those detected by other methods, although MTClass shared more top genes with MultiPhen and MANOVA than with single-tissue eQTL and sQTL methods (**Figures 3C** and **3D**).

**Figure 3.**
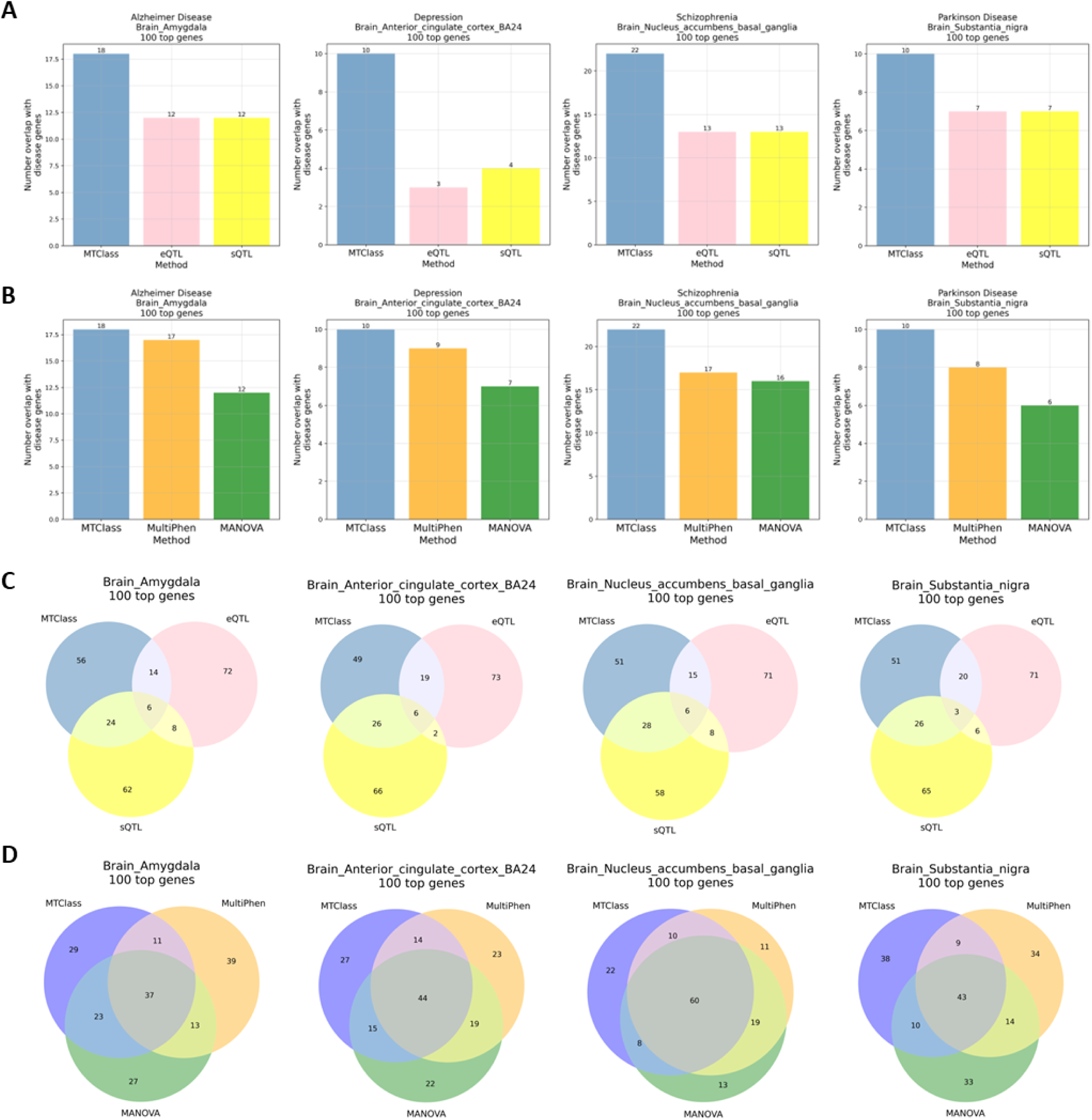
Functional assessment of top genes identified by MTClass compared to single-tissue studies and other multivariate methods in the multi-exon study in the brain. All overlap analyses were conducted using the DisGeNET database, version 3. **A**. Overlap with known disease-associated genes among the top 100 genes compared to single-tissue eQTL and single-tissue sQTL methods. **B.** Overlap with known disease-associated genes among the top 100 genes compared to MultiPhen and MANOVA. **C.** Overlap among the top 100 genes detected by single-tissue methods. MTClass largely detects distinct top genes from single-tissue eQTL and sQTL methods. **D.** Overlap among the top 100 genes detected by MTClass compared to MultiPhen and MANOVA.

### Multi-tissue and multi-exon 2D study

Given the highly correlated nature of gene expression among exons and tissues, a third consideration is that one eQTL can have far-reaching effects across both exons and tissues. We tested this possibility by using a two-dimensional feature matrix of multiple (m ≥ 10) exons across nine tissues for each gene as input to our model. We used the same tissues as in the 9-tissue case to avoid imputing missing data. The model we chose for this study was a MLP, or fully connected neural network, to facilitate the capture of any latent structure among the expression levels.

We also tried the same data (flattened vector) on MultiPhen and MANOVA; however, these two methods encountered multiple numerical issues due to the high dimensionality and abnormal correlation pattern of the data. Thus, we decided not to pursue a performance comparison on this dataset. We ran only one iteration of MTClass on this dataset due to the relatively higher computational burden. The top-performing eQTLs of the top 10 eGenes ranked by MCC from the 2D study are summarized in **Table 3**.

One eGene-eQTL pair that achieved a perfect macro F1 score and MCC in the 2D study was FAM118A-chr22_45315973_G_A_b38 (rs104664). This SNP is located in the intronic region of the FAM118A gene on chromosome 22. It is an eQTL in 49 out of 54 tissues (90.7%), including all 9 tissues used in this study. Intriguingly, this SNP is also an sQTL in 50 out of 54 tissues (92.6%), including all 9 tissues used in this study. When examining the normalized gene expression between genotypes, it is evident that the two genotypes have markedly different expression patterns across multiple exons and multiple tissues (**Figure 4A**). Upon nonlinear dimensionality reduction via UMAP, the genotypes are clearly highly separable (**Figure 4B**), providing a possible explanation for our method’s ability to classify this SNP well. This SNP has been reported as a significant eQTL in human osteoblasts^50^, sputum^51^, as well as CD4+ lymphocytes^52^, suggesting that it has complex, cross-tissue effects on gene expression within the body. Furthermore, FAM118A is an integral membrane protein, and although its exact functions remain unclear, it is highly expressed in many regions of the body, suggesting it has important functions across several somatic tissues.

**Figure 4.**
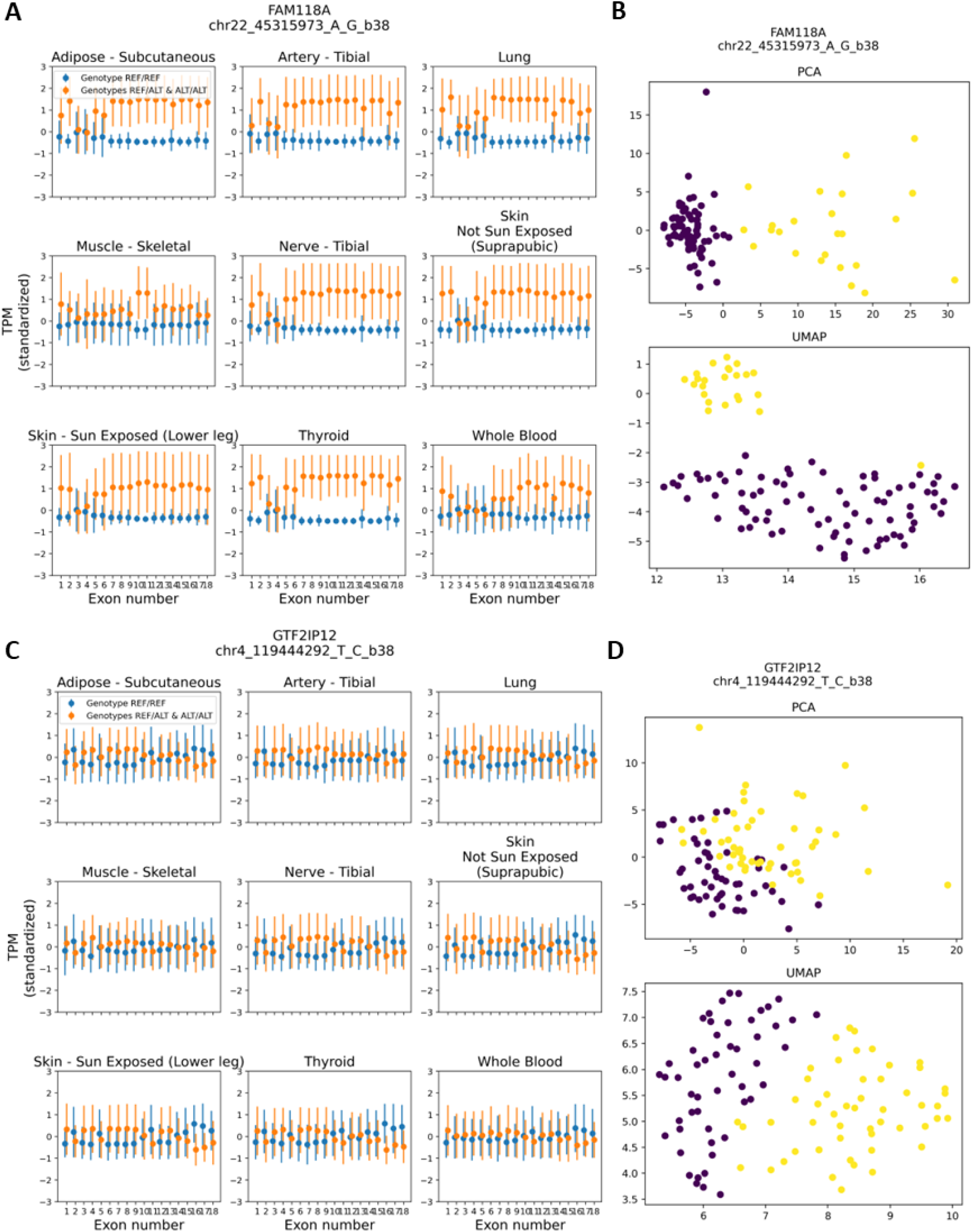
Anecdotal examples of eGene-eQTL pairs classified well by MTClass across multiple exons and tissues in the 2D study. **A.** Mean and standard deviation plots of standardized expression levels in TPM across all exons and 9 tissues. This eGene-eQTL pair shows differences in gene expression in multiple exons and multiple tissues. **B.** Both PCA and UMAP dimensionality reduction show high separability between the two genotypes. **C.** Expression levels in TPM across exons and tissues for an eGene-eQTL pair exhibiting less separability between genotypes. **D.** Linear dimensionality reduction with PCA cannot separate the genotypes as neatly as nonlinear techniques such as UMAP, suggesting the relationship is more nonlinear.

Furthermore, an eQTL that exhibits a more nonlinear relationship between genotype and multiple phenotypes is GTF2IP12-chr4_119444292_T_C_b38 (rs9759690). This eQTL achieved a perfect macro F1 score and MCC. According to the GTEx Portal, this is a significant eQTL in 36 out of 54 tissues (66.7%), including six out of the nine tissues (66.7%) used in this study. It is also a significant sQTL in 19 out of 54 tissues (35.2%), including six out of the nine tissues (66.7%) used in this study, demonstrating that it is indeed a multi-tissue, multi-exon eQTL. **Figure 4C** shows that the relationship between genotype and phenotype is complex. Intriguingly, principal component analysis (PCA) was less able to separate the genotypes compared to UMAP, a nonlinear dimension reduction technique (**Figure 4D**). This suggests that the genotype-phenotype relationship is probably more nonlinear and therefore better captured by MTClass.

### PsychENCODE isoQTL study

In vertebrate eukaryotic species, alternative splicing generates an enormous amount of mRNA diversity by facilitating the formation of multiple mRNA isoforms. Isoform diversity is thought to contribute to a cell’s ability to adapt to the cellular microenvironment. Aberrant alternative splicing has been shown to contribute to neurological disorders such as AD and PD^53,54^. Thus, understanding the contribution of genetic variation to the abundance of various mRNA isoforms will provide a more comprehensive understanding of alternative splicing and neurological illness.

Because of the relatively small sample sizes available in the 9-tissue and brain tissue studies (*n* = 103 and *n* = 317, respectively) from the GTEx consortium, we turned to an independent dataset from the PsychENCODE Consortium^19,21^, specifically the subset of data from the CommonMind Consortium (CMC)^20^. Here, each feature was the expression level in TPM from each isoform of a gene, obtained from post-mortem adult prefrontal cortex of the brain. We restricted our analyses to genes that had at least 10 isoforms to ensure there were enough features for our algorithm to consider. There was a total of *n* = 900 donors in this study. We term the top-scoring variants “isoQTLs” and their corresponding genes “isoGenes” because the variants are associated with the expression of multiple isoforms of a gene.

We ran MTClass three times on the same dataset with different random seeds. The hyperparameters were kept the same. MultiPhen and MANOVA were also run as a comparison. Subsequently, we conducted GWAS variant colocalization analysis on the top 5,000 variants for each method. We found no marked differences in GWAS variant colocalization among the top variants detected by the three methods. Secondly, like our analysis in the multi-exon studies in brain tissue, we calculated the overlap with known disease-related genes retrieved from DisGeNET. We found that MTClass identified more isoGenes associated with anxiety disorder and bipolar disorder, two diseases that significantly affect prefrontal cortex function^55–58^ (**Figure 5A**). Moreover, we observed that for other prefrontal cortex-related disorders such as schizophrenia, autism spectrum disorder, attention-deficit hyperactivity disorder, and obsessive-compulsive disorder, MTClass always achieved as much, or even greater, disease-gene overlap as MultiPhen and MANOVA **(Supplementary Figure 9)**. The top-performing isoQTLs of the top 10 isoGenes ranked by MCC are summarized in **Table 3**. Altogether, these results suggest that MTClass excels in recapitulating functionally relevant genes in the prefrontal cortex.

**Figure 5.**
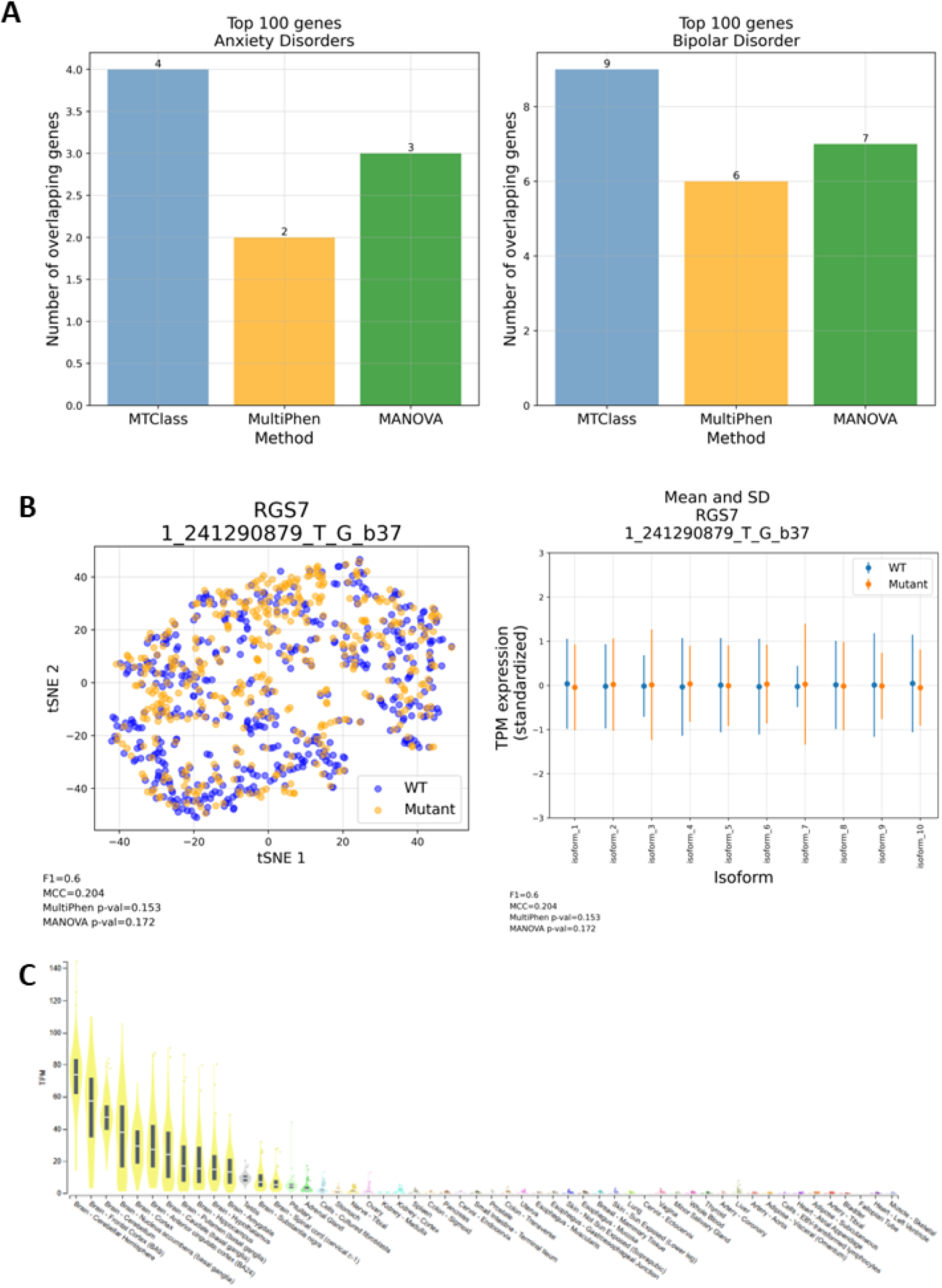
Functional assessment of variants and genes detected by MTClass compared to other methods in the isoQTL study in human prefrontal cortex. **A.** Using the DisGeNET database version 3, top isoGenes detected by MTClass show greater overlap with prefrontal cortex-associated disorders like anxiety disorder (left) and bipolar disorder (right) compared to those from other methods. **B.** Anecdotal example of an isoGene-isoQTL pair detected by MTClass but not by other methods. t-SNE dimensionality reduction (left) of this eQTL shows poor separation of genotypes, but suggestive regions where one genotype predominates. This isoGene shows expression differences across multiple isoforms and differences in variance in isoforms 3 and 7, where the mutant genotype has higher variation compared to the wild type genotype (right). **C.** According to the GTEx Consortium, the RGS7 gene is most highly expressed in the brain, especially in the frontal cortex.

We provide an anecdotal example of an isoGene-isoQTL pair (RGS7-1_241290879_T_G_b37 (rs10926417)) that was classified well by MTClass but did not achieve statistical significance in either MultiPhen or MANOVA. MTClass achieved a reasonably high median macro F1 score of 0.60 and a median MCC of 0.204, while MultiPhen reported a p-value of 0.153, and MANOVA reported a p-value of 0.172.

Dimensionality reduction using t-SNE suggests that the two genotypes are tightly intertwined, with a few regions where one genotype predominates. However, examining the isoforms separately, it is evident that while the mean expression levels are similar across isoforms, the variance of the mutant genotype in isoforms 3 and 7 is significantly higher than that of the wild-type genotype (**Figure 5B**). This suggests a potential dysregulation of the expression of these specific RGS7 isoforms in individuals with the mutant genotype. Importantly, RGS7 is highly abundant in the brain, with the highest median expression in the cerebellar hemisphere and the frontal cortex (**Figure 5C**). As a regulator of G-protein signaling, RGS7 has been linked to intellectual disability and depression-related behaviors^59,60^.

### OneK1K scRNA-seq study

Most eQTL studies have been conducted on bulk RNA-sequencing data, in which the data represent an average expression of a gene across all cells in a tissue^61^. The advent of single-cell RNA-sequencing (scRNA-seq) has revolutionized the field of transcriptomics by allowing the quantification of cell-type-specific gene expression and the identification of cell type heterogeneity^62–65^. Cell-type-specific eQTLs have been identified in scRNA-seq studies as well, although this focuses on a single cell type at a time^22,66^. It is likely that some eQTLs can have effects on more than one cell type. Thus, we extended our study to identify eQTLs that affect gene expression level vectors across multiple cell types.

The OneK1K cohort is a notable example of the power of scRNA-seq, sequencing over a million cells from peripheral blood mononuclear cells (PBMCs) across nearly 1,000 individuals^67^. We obtained this dataset from The Human Cell Atlas. We chose the seven major cell types that contained at least one cell for all 982 donors, resulting in no missing cell-type/donor combinations (**Figure 6A**).

**Figure 6.**
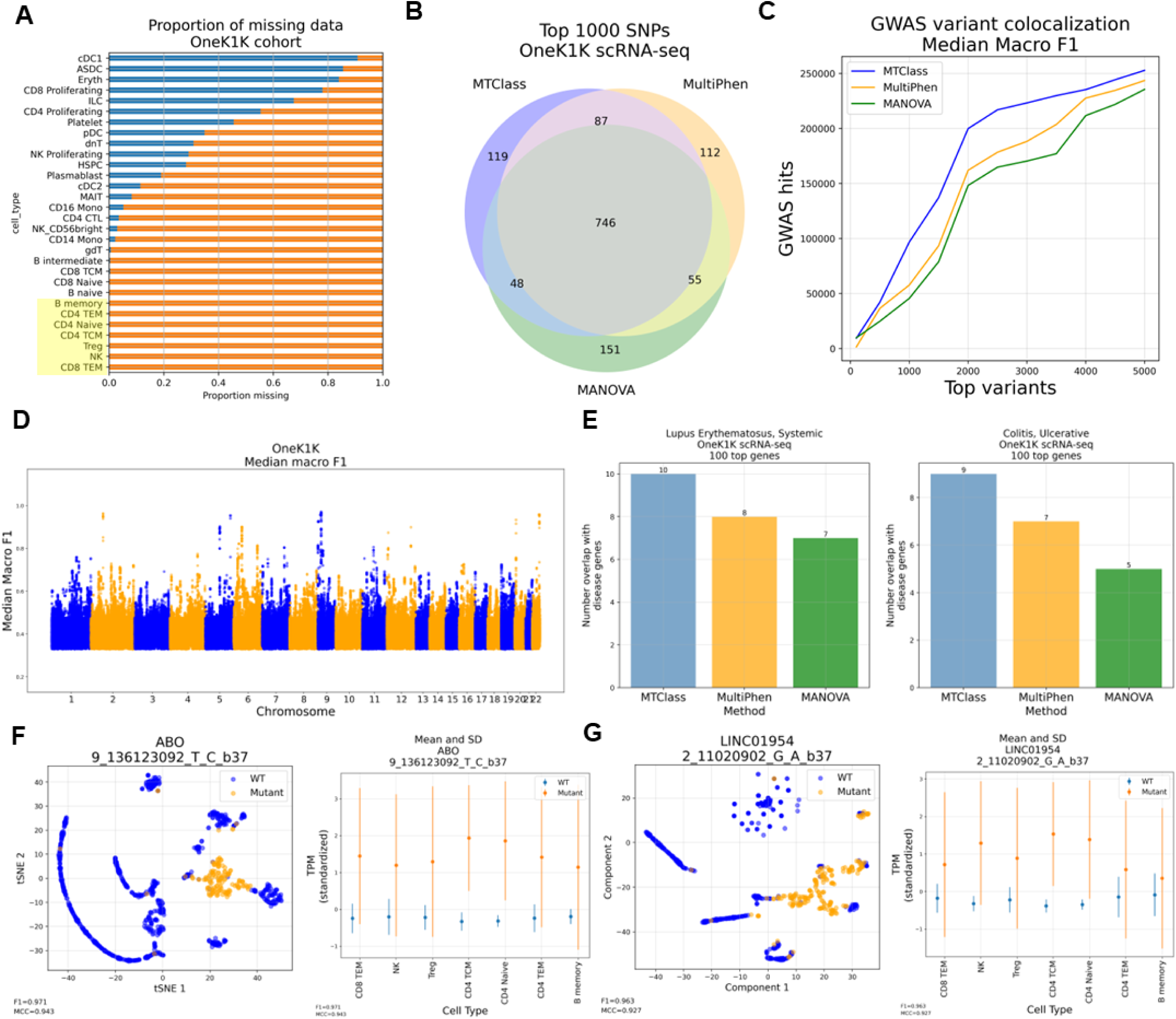
MTClass identifies eQTLs across multiple cell types in the OneK1K scRNA-seq cohort. **A.** Visualization of missing data across different cell types in the dataset. The yellow highlighted box indicates the 7 cell types that were used as features for multi-cell-type eQTL identification. **B.** There is a substantial amount of overlap in the top 1000 variants detected by each multivariate association method. **C.** MTClass identifies variants that colocalize with more known trait-associated SNPs from the GWAS Catalog compared to MultiPhen and MANOVA. **D.** Manhattan plot of MTClass results across 3 iterations by median macro F1 score. **E.** The top MTClass genes overlap with more genes associated with various autoimmune diseases such as systemic lupus erythematosus and ulcerative colitis compared to the top genes from MultiPhen and MANOVA. **F.** The top MTClass result is a SNP in the ABO gene, which helps to determine human blood typing. UMAP analysis shows highly separable genotypes (left). The mutant genotype has both higher expression and higher variance in this gene compared to the wild type genotype in all seven cell types (right). **G.** LINC01954-2_11020902_G_A_b37 (rs7577665) is an eQTL-eGene pair among MTClass’ top results that is not a known eQTL in whole blood from the GTEx Consortium data. **A.** t-SNE analysis clearly shows that the genotypes are separable (left). Across several cell types, the mutant genotype has both a higher mean expression and higher variability in expression levels (right).

We ran our MTClass classifier on the OneK1K dataset, using expression measures from the seven major cell types as features for each gene, and we compared our approach to MultiPhen and MANOVA. We observed that there was a substantial amount of overlap among the top 100 genes detected by each of the three multivariate association methods (**Figure 6B**), which could be due to the highly correlated nature among the seven selected cell types in the OneK1K cohort. After conducting the GWAS variant colocalization analysis, we observed that the top variants identified by MTClass colocalized with more known signals in the GWAS Catalog compared to those identified by MultiPhen and MANOVA (**Figure 6C**). We also noticed that most of these signals originated from chromosome 6, on which many HLA genes reside **(Supplementary Figure 10)**.

Figure 6D depicts the Manhattan plots for MTClass based on median macro F1 score and median MCC, corroborating the observation that there are strong signals from chromosome 6. This evidence suggests that immune-related traits comprise most of the identified signals, which is biologically plausible because most of the seven cell types included in our study have important functions in immune signaling and antigen processing.

Using the DisGeNET database, we found that the top eGenes selected by MTClass overlapped with more genes associated with various autoimmune disorders such as systemic lupus erythematosus and ulcerative colitis (Figure 6E).

The top-performing eQTLs of the top 10 eGenes ranked by MCC from the OneK1K scRNA-seq study are summarized in **Table 3**. One eQTL that was detected robustly by MTClass was ABO-9_136123092_T_C_b37 (rs11244049), which is a downstream gene variant in the ABO gene on chromosome 9. This SNP achieved a median macro F1 score of 0.971 and a median MCC of 0.943. t-SNE dimensionality reduction analysis was able to segregate the genotypes quite well. This SNP also seems to have a pronounced effect on ABO gene expression, with the mutant genotype having markedly higher mean expression and a greater variance compared to the wild-type genotype (Figure 6F). Unsurprisingly, many of the top variants identified by MTClass modulate the ABO gene, which is responsible for human blood typing^68^. This result makes intuitive sense given that the data are from human peripheral blood.

Among the top 10 MTClass results listed in **Table 3**, LINC01954-2_11020902_G_A_b37 (rs7577665) is an eGene-eQTL pair that is not a known pair in whole blood on the GTEx Portal. According to the GTEx Consortium, this pair is only statistically significant in the spleen. MTClass was able to classify this pair with high precision (macro F1 score = 0.963, MCC = 0.927). Indeed, dimensionality reduction using t-SNE shows that the genotypes are clearly separable. Across the seven cell types, the mutant genotype generally exhibits higher mean expression levels as well as greater variation in expression compared to the wild-type genotype (Figure 6G). Altogether, we show that MTClass is a powerful and effective tool to detect eQTLs across cell types in scRNA-seq studies.

### Comprehensive performance comparison

Several other statistical methods have been developed to perform multivariate genetic association. In this study, we consistently compared the performance of MTClass to MultiPhen^11^ and MANOVA^12^ to alleviate the computational burden of running all methods on all datasets. However, for the brain tissue case, we compared MTClass to a diverse array of methods to illustrate the superior performance of our approach.

Notably, we utilized canonical correlation analysis (CCA, as implemented in mv-PLINK^69^), multivariate logistic regression (an adaptation of SCOPA/META-SCOPA^70^), and the Cauchy combination test^71^ to combine p-values from single-tissue eQTL association studies. Specifically, we used GWAS variant colocalization analysis to evaluate the potential of each method at detecting functionally relevant eQTLs. We observed that MTClass outperformed all other methods on this metric in the brain tissue case (**Supplementary Figure 11**), demonstrating that MTClass excels at detecting genetic variants with high functional importance.

## Discussion

Understanding how genetic variants influence transcription regulation is highly important. Given that the transcription process is tightly regulated, it is of little surprise to find that genetic variants, *i.e.* eQTLs, can affect gene expression across multiple entities. Therefore, the identification and understanding of such eQTLs with pleiotropic effects on gene expression is of particular interest. In this study, we proposed to use machine learning algorithms to classify donor genotypes based on the vector of expression levels from multiple sources (e.g., tissues, exons, isoforms, and cell types). To the best of our knowledge, this is the first time that a classification-based approach was utilized to identify eQTLs. We compared our approach to other avant-garde linear approaches, namely MultiPhen and MANOVA, and showed that our method often detects eQTLs and eGenes with greater functional impact than other methods. In this way, we introduce a novel means of conducting multivariate association analyses, which could motivate the development of future multivariate trait association methods.

We conducted simulation studies to demonstrate the utility of our approach in uncovering nonlinear associations between genotypes and phenotypes. In two hypothetical examples using three phenotypes, we showed that MultiPhen and MANOVA failed to detect the nonlinear association, whereas MTClass classified the simulated “genotype” almost perfectly. Admittedly, the simulated extreme nonlinear patterns of genotype-phenotype associations may be rare in real data, but we hypothesized that nonlinear associations are likely to be ubiquitous nevertheless, especially when considering a large number of phenotypes.

We next tested MTClass on multiple real datasets. In most cases, we demonstrated that MTClass outperformed MultiPhen in detecting eQTLs of functional importance especially in the brain tissue case. Upon examining the top variants, we observed that variants associated with the HLA genes on chromosome 6 contributed significantly to the increase in GWAS hits in the brain tissue case. These variants tend to be classified very well by MTClass, but not by MultiPhen or MANOVA, suggesting a more nonlinear relationship between genotype and phenotype. Thus, many immune-related variants and genes are exclusively detected by MTClass, suggesting that such immune-related variants and genes exhibit complex pleiotropic effects across multiple somatic tissues. Consequently, we hypothesized that existing approaches have underestimated the number of eQTLs and their complex effects on HLA genes in the MHC region, which play essential roles in immune function and immune-related diseases. Nevertheless, we acknowledge that eQTL mapping in the MHC region is extremely difficult due to the complexity of the region^72^. Future work should take advantage of whole-genome sequencing to perform eQTL mapping specifically in the MHC region.

In the multi-exon study, we observed that the top eGenes detected by MTClass overlapped more with known disease-associated genes retrieved from DisGeNET. We calculated the number of overlapping genes to demonstrate that MTClass detects eGenes with greater functional importance compared to single-tissue and other multivariate methods. As expected, we also found that all three tested multivariate approaches (MTClass, MultiPhen, and MANOVA) recovered a greater number of known disease genes compared to the single-tissue eQTL and sQTL approaches in brain tissue. This suggests that disease-associated genes are more likely to be regulated by pleiotropy and are consequently better captured by methods that consider more than one phenotype.

Lastly, we demonstrated in the 2D multi-exon, multi-tissue study that our MLP-based algorithm was able to detect variants that were significant as both eQTLs and sQTLs in multiple tissues, thereby contributing broadly to both gene expression and splicing events. We also provided an anecdotal example of an eGene-eQTL pair that exhibited a more nonlinear association, which could not be captured by MultiPhen or MANOVA.

Compared to existing methods, MTClass has the following advantages. First, it provides a generalized framework to conduct association studies. As such, single-tissue eQTL mapping is a special instance of our framework in which there is only one feature. Even in the multivariate case, if an eQTL significantly affects gene expression in only one phenotype, MTClass would still classify it well. Second, our results demonstrated that MTClass identifies more functionally important eGenes and eQTLs than competing methods which are linear model-based, suggesting that classification-based methods are better at identifying non-linear patterns of separation. Third, multicollinearity between features poses less of a problem for MTClass, as the algorithm can learn the redundancy of a set of features through training. In contrast, linear multivariate methods such as MultiPhen and MANOVA will often fail when two independent variables are collinear. Lastly, MTClass is more flexible as it accepts a greater range of values as input, including features with zero expression in all donors. An example of such a feature can be found in exon-level expression, in which certain exons might not be expressed at all in certain genes due to exon skipping during alternative splicing. Conversely, MultiPhen and MANOVA will both report numerical errors if an independent variable is zero across all donors. Therefore, we believe that MTClass is a more reliable and flexible approach compared to linear multivariate methods such as MultiPhen and MANOVA.

One limitation of this study was the relatively low sample size in the 9-tissue case and in the 2D exon-tissue case, where expression levels and genotypes were available for only 103 donors. This would have a much larger negative impact on our results than on those of MultiPhen and MANOVA, since machine learning-based methods make no modeling assumption and in return, require a large sample size to accurately detect patterns in the data. We utilized an independent dataset of human prefrontal cortex from the PsychENCODE Consortium and a scRNA-seq dataset to alleviate this issue and validate our GTEx results. Indeed, we showed that in the PsychENCODE isoQTL study, MTClass detected genes that overlapped with more prefrontal cortex-related phenotypes. Moreover, we provided evidence that in the OneK1K cohort with 982 samples, GWAS variant colocalization analysis favored MTClass over both MultiPhen and MANOVA. We also concede that multiple imputation is better suited to data missing at random, not systematically missing data such as in the GTEx samples. Further investigation needs to be conducted into how to impute cross-tissue gene expression with greater accuracy.

Another limitation is that we only considered two groups of genotypes by combining heterozygous and homozygous mutants into one, in accordance with a dominant model for genetic association studies. We did this due to the limitation of sample size and of the lower minor allele frequency. By combining these two genotype classes, we alleviated the problem of class imbalance, which is known to negatively impact model performance, especially in cases of lower sample size. In doing so, we assumed that the heterozygous and homozygous mutants carried similar genetic risk. We acknowledge that this assumption might not be valid for some variants and that using three distinct genotype classes might be more appropriate in such cases^73^.

In traditional eQTL mapping, covariates such as age, sex, and genotyping principal components are routinely incorporated as potential confounders in the linear models employed. Since it is challenging to include these covariates under a classification framework, we believe this is an important consideration for further development of our method to filter out potential false positives.

So far in genetics, all association tests are conducted under the hypothesis testing framework. Despite overwhelming success, this strategy does not work well for multivariate traits. In a recent study, Yu et al. successfully applied classification-based GWAS to full brain magnetic resonance (MR) images^74^. In this study, we showed that a similar strategy can be applied to quantitative traits such as eQTLs. To the best of our knowledge, this is the first time that a classification-based approach was utilized to identify eQTLs.

Overall, identifying multi-phenotype eQTLs has the potential to elucidate biologically meaningful effects of genotype on gene expression and to provide insights into pleiotropic interactions. We are certainly not the first group to propose that quantitative trait loci could affect disparate phenotypes such as gene expression, alternative splicing, and even polyadenylation^75^. Our results will need to be validated in other datasets and experimentally, but we believe they provide an exciting new avenue for conducting multivariate association analyses using machine learning.

## Data availability

Classification metrics from MTClass, as well as association p-values from MultiPhen and MANOVA, for the multi-tissue study (9-tissue, 13 brain tissues, 48-tissue), the multi-exon study in 13 brain tissues, the multi-tissue/multi-exon 2D study, the isoQTL study using PsychENCODE data, and the OneK1K scRNA-seq study are available via Zenodo at https://doi.org/10.5281/zenodo.10911584. The original GTEx data used for this study can be found at https://gtexportal.org/, and the subset of data from the PsychENCODE CommonMind Consortium can be found via Synapse at https://www.synapse.org/#!Synapse:syn12080241. The OneK1K single-cell gene expression and genotype data are available via Gene Expression Omnibus (GSE196830). The cell by gene matrix is available at The Human Cell Atlas (HCA) (https://cellxgene.cziscience.com/collections/dde06e0f-ab3b-46be-96a2-a8082383c4a1).

## Code availability

The MTClass algorithm implements standardization of expression measures, model training, and classification using an ensemble of random forest and support vector machine. One of the evaluation metrics used to assess the functional relevance of variants was GWAS variant colocalization. The source code, along with example data, is available via GitHub at https://github.com/ronnieli0114/MTClass.

## Methods

### Selection of genes and variants

To prioritize genes with diverse expression levels, we calculated the expression variance for all genes across all 17,382 tissue samples (across donors) and excluded genes that had expression levels of zero in more than half of these samples. We then selected the top 50% of genes (N = 13,810) in terms of variance.

For each of the selected variable genes, we used PLINK to extract all of its candidate *cis*-eQTLs within 10 kb of the gene’s coding region, and only retained variants that have a minor allele frequency (MAF) of at least 0.05. In accordance with a dominant genetic model, the homozygous reference genotype was denoted as 0, while the heterozygous and homozygous mutant genotypes were denoted as 1. We combine genotypes that contain at least one mutant allele to ensure that there are enough samples with the least common genotype category for the machine learning algorithm to learn from the data. We removed variants and donors with a genotyping rate less than 90%, and we restricted our analysis only to autosomes to avoid potential issues with X-chromosome haploinsufficiency affecting expression levels.

### Imputation of gene expression measures

To impute missing expression measures for the brain tissue case (K = 13 tissues) and the 48-tissue case (*K* = 48 tissues), we tested a variety of multiple imputation techniques. Multiple imputation iteratively treats each feature (*i.e.*, tissue) as the response variable in a linear regression, using a combination of the known values of other features to predict the missing values^32,76^.

We randomly masked 50% of the 927 samples for the 9-tissue case to test the imputation strategies. The pattern of missing data was the same as the real GTEx data in the brain tissue and 48-tissue cases. For each gene, we measured the Pearson correlation coefficient (*R*^2^) between the predicted values and the actual values. We tested five multiple imputation techniques: (1) support vector regression^77^, (2) Bayesian ridge regression^78^, (3) random forest regression^79^, (4) predictive mean matching^80^, and (5) K-nearest neighbors regression^81^. For all methods except predictive mean matching, we employed the IterativeImputer() object in the Python scikit-learn package with 30 maximum iterations. For predictive mean matching, we used the MICE() object with five iterations from the Python statsmodels package to perform the imputation. Overall, we found that predictive mean matching yielded the highest median R^2^, so we imputed the expression levels for the brain tissue case using this method **(Supplementary** Figure 5).

### Model selection and evaluation

Figure 1 depicts a schematic overview of the MTClass algorithm. We chose to utilize an ensemble approach comprised of random forest and support vector machine as base learners for our machine learning model. We found that the ensemble of these two base learners reduced overfitting and improved overall performance. Specifically, we used 150 estimators for the random forest approach, and we used a radial basis function (RBF) kernel for the support vector machine with C = 1.0. Hyperparameters were decided based on a grid search algorithm. The RBF kernel is the most conducive to capturing nonlinear relationships, as opposed to others like the linear kernel. We employed a soft-voting approach, where the predicted class for each sample was the *argmax* of the summed probabilities of each base classifier. The ensemble classifier was implemented using the Python scikit-learn package.

In the multi-exon/multi-tissue 2D study, because our feature matrix was two-dimensional, we used a MLP with one hidden layer instead of an ensemble algorithm. This ensured that our network would be capable of detecting any latent structure among the features. The MLP was implemented in the Python Tensorflow/Keras package. We considered the fact that each gene would have a different number of features, namely *m* × *n*, where *m* is the number of exons and *n* is the number of tissues. Therefore, the size of the single hidden layer was changed for each gene to be ^*mn*^, rounded up to the nearest integer.

We combined either the ensemble approach or the neural network approach with 4-fold stratified cross validation to mitigate model overfitting. We excluded variants that had fewer than 4 samples in the less common class so that each stratified fold would have at least one sample with the less common class. Finally, we standardized each feature using the z-score method immediately prior to fitting our classifier. This procedure guaranteed that all the features would be on the same scale, which is critical for preventing bias in the machine learning model.

To evaluate classification performance, we used the macro F1 score and MCC. Both metrics are robust to class imbalance, which was essential to our choice, as some eGene-eQTL pairs contained few samples of the less common class.

### Simulation study

In the first simulation experiment, we generated 2 groups of 100 data points each with the same arithmetic mean but different variances in a spherical pattern. One group of 100 samples had the “0” genotype, while the other 100 samples had the “1” genotype. To easily visualize this data, we only used 3 “tissues,” or features. We randomly generated two angles, θ and ø, 0 ≤ θ < 2π and 0 ≤ ø < π, such that the features (*x, y, z*) were given by:

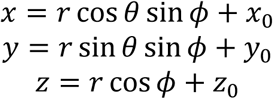

Here, *r* denotes the radius of the sphere. For one sphere, *r* was set to be 20, and for the other, *r* was set to be 5. Thus, the two groups had the same mean (*x*_0_, *y*_0_, *z*_0_), but different variances.

In the second experiment, we still generated 2 groups of 100 data points each, but in interlocking sinusoidal patterns. We continued to use 3 features to facilitate visualization. We randomly generated features (*x, y, z*) such that:

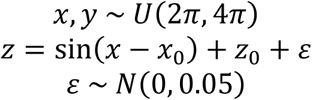

Here, *x* and *y* were sampled from a uniform distribution, and ε denotes Gaussian noise added to introduce additional complexity to the dataset.

### Comparison with other methods

We compared our machine learning method with two state-of-the-art multivariate association methods, MultiPhen and MANOVA. Both methods output a single nominal p-value such that the ranking of the SNPs is possible for the purpose of performance evaluation. For this reason, we did not include methods such as MASH^14^.

MultiPhen^11^ performs a “reverse” ordinal regression on the genotypes to test for the significance of association with multiple phenotypes. Here, the variant genotypes become the dependent variable, and the *K* phenotypes become the predictor variables. Proportional odds logistic regression is used to define the probabilities for each class (allele count):

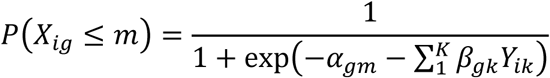

We used the mPhen() function in the MultiPhen R package with default parameters to run the analysis.

MultiPhen is installable from CRAN (http://cran.r-project.org/) or any CRAN mirror, with documentation available at http://cran.r-project.org/web/packages/MultiPhen/MultiPhen.pdf.

Multivariate analysis of variance (MANOVA)^82^ is a statistical test commonly used to test the significance of association between a predictor variable (genotype) and several response variables (phenotypes). MANOVA assumes that the phenotypes are normally distributed and that there is no multicollinearity between the dependent variables. We used the built-in manova() function in the R language with default parameters to conduct the analysis. The dependent variables were the *K* phenotypes, and the independent variables were the variant genotypes. By default, the manova() function in R tests the significance of the Pillai trace statistic.

Because MultiPhen and MANOVA sometimes reported numerical errors for certain eGene-eQTL pairs, we excluded features with expression levels of zero in all donors, allowing MultiPhen and MANOVA to conduct the association for a greater number of eGene-eQTL pairs. Lastly, we ensured all methods tested on the same gene-variant pairs, and we excluded all pairs for which either MultiPhen or MANOVA reported a numerical error.

In the brain tissue case, we also compared the performance of MTClass to that of three additional methods: CCA, reverse logistic regression, and the Cauchy combination test. CCA identifies and measures the associations among two sets of variables, and it is especially useful when there are multiple intercorrelated variables such as phenotypes. CCA has been implemented in mv-PLINK^69^, but because we did not have access to the software, we implemented a modified version using the cancor() function in R. We then tested the statistical significance of the correlation value ρ using the Wilks’ lambda statistic in the CCP package in R.

Akin to MultiPhen, SCOPA and META-SCOPA^70^ utilize a reverse linear regression in which the genotype is the outcome variable and the phenotypes are the predictor variables. However, due to the unavailability of the software, we implemented a modified version of the model but used logistic regression to more effectively represent the binarized genotypes. Specifically, we compared the full model, in which all phenotypes were used as predictors, to a null model in which no predictors were used. We used analysis of variance to calculate a p-value quantifying the improvement in model fit achieved by adding the predictors.

Finally, we calculated nominal p-values for each eGene-eQTL pair in each of the 13 individual brain tissues using tensorQTL^83^. We used all covariates computed by the GTEx Consortium version 8 in the analysis. Next, we combined the 13 p-values using the Cauchy combination test^71^ to obtain a single p-value for each eGene-eQTL pair. The Cauchy combination test was implemented in the ACAT package in R.

### GWAS variant colocalization analysis

GWAS variant colocalization analysis was done to assess the relative functional importance of the top variants identified by each method. We downloaded the entire GWAS Catalog^26^ from the NHGRI-EBI website: https://www.ebi.ac.uk/gwas/. Entries with duplicate or missing genomic positions were removed.

Because the GWAS Catalog uses the hg38 reference genome, we utilized the UCSC liftOver tool (https://genome.ucsc.edu/cgi-bin/hgLiftOver) to convert the coordinates from hg38 to hg19 to be compatible with the genotype data from the PsychENCODE isoQTL and OneK1K scRNA-seq studies. In doing so, 138 coordinates could not be converted and were excluded from the analysis. The total number of unique entries left in the GWAS Catalog that contained both hg38 and hg19 genomic coordinates was 536,898.

We sorted the MTClass results by classification metric (macro F1 score or MCC) in descending order, and we sorted the results from all other methods by p-value in ascending order. Then, we selected the top *N* variants from the sorted results, and for each top variant, we computed the number of unique genomic positions in the GWAS Catalog that were within 10 kilobases of the variant’s genomic locus. We termed this number the “GWAS hits.”

When selecting the top *N* variants, it is possible that we are unable to select exactly *N* top variants due to ties in p-value or classification performance. To account for this, for each result, suppose the fewest number of variants that produce at least *N* top variants is *M*. Therefore, *M* ≥ *N*. We next calculated the number of GWAS hits for these *M* variants, then adjusted this count by *N*/*M*.

### Disease gene overlap analysis

DisGeNET (https://www.disgenet.com) hosts one of the largest publicly available collections of genes and variants associated with human diseases^49^. We first downloaded disease-gene associations for various neurological diseases from version 3.0 of the database. For each study, we filtered out disease genes that were not in the background gene list. Next, we selected the top 100 unique eGenes from the MTClass results after sorting by median macro F1 score in descending order. We first compared the MTClass result to single-tissue results by downloading the single-tissue eQTL and sQTL datasets from the GTEx Consortium. We sorted their eGenes by nominal p-value in ascending order, then selected the top 100 unique eGenes. We did the same with MultiPhen and MANOVA after sorting by nominal p-value in ascending order. Finally, we computed the intersection of the result sets with the filtered disease gene list to obtain the overlap.

### OneK1K scRNA-seq data processing

This study aimed to identify eQTLs across multiple cell types from single-cell RNA-sequencing data. We used peripheral blood mononuclear cells (PBMCs) sequenced in the OneK1K cohort due to the large sample size. Raw scRNA-seq data were downloaded from CZ CELLxGENE Discover. The data were processed using the Seurat v4 and dplyr packages in R. First, the raw Unique Molecular Identifier (UMI) counts were normalized with the NormalizeData() function in Seurat. Default parameters were used, resulting in a standard log-normalization procedure. Then, the FindVariableFeatures() function was used to find and retain the 3,000 genes with the highest variance. We examined the number of donors available for each annotated cell type, and we kept the 7 cell types with all 982 donors, resulting in no missing data. These 7 cell types constituted approximately 77% of all predicted cells in the dataset, including the 4 most abundant cell types. Specifically, the chosen cell types were B memory, CD4 TEM, CD4 Naïve, CD4 TCM, T regulatory, natural killer (NK), and CD8 TEM. To construct the feature matrix for our classifier, for each gene, the sum of the normalized expression levels of all cells (pseudobulk) belonging to a given cell type was taken as the expression measurement for that cell type. The features were finally z-standardized using Python’s StandardScaler() object before running the MTClass algorithm.

The genotype data for the OneK1K cohort were obtained via personal communication and converted to PLINK format for efficient access. We followed the same procedure to extract *cis*-eQTLs of interest as in the previous experiments. We compared our approach to MultiPhen and MANOVA. All methods were run on binarized genotypes in accordance with a dominant model.

### Variant annotation

For each of the MTClass results, the top eQTLs by MCC from the top 10 eGenes were annotated using Ensembl’s Variant Effect Predictor (VEP, release 113)^84^: https://useast.ensembl.org/info/docs/tools/vep/index.html. The GRCh37 version of the VEP was used for the isoQTL and OneK1K scRNA-seq results, while the GRCh38.p14 assembly was used for all others. To resolve duplicate annotations, only the most severe functional consequence and one rsID was kept for each variant.

## Author contributions

All authors contributed extensively to this work. R.Y.L. and Z.S.Q. conceived the idea for the MTClass algorithm. R.Y.L. assembled input data, wrote the code, analyzed the output data, and wrote the draft of the manuscript. C.S. and Z.S.Q. supervised the study and made substantive revisions to the manuscript.

## Ethics statement

The authors declare no competing interests.

## Materials and correspondence

Correspondence should be addressed to Dr. Zhaohui “Steve” Qin at.

## Biographical note

Ronnie Li is an alumnus of the Neuroscience Graduate Program at Emory University. Chang Su and Zhaohui Qin are faculty members at Emory University

## Supplementary Figures

**Supplementary Figure 1.**
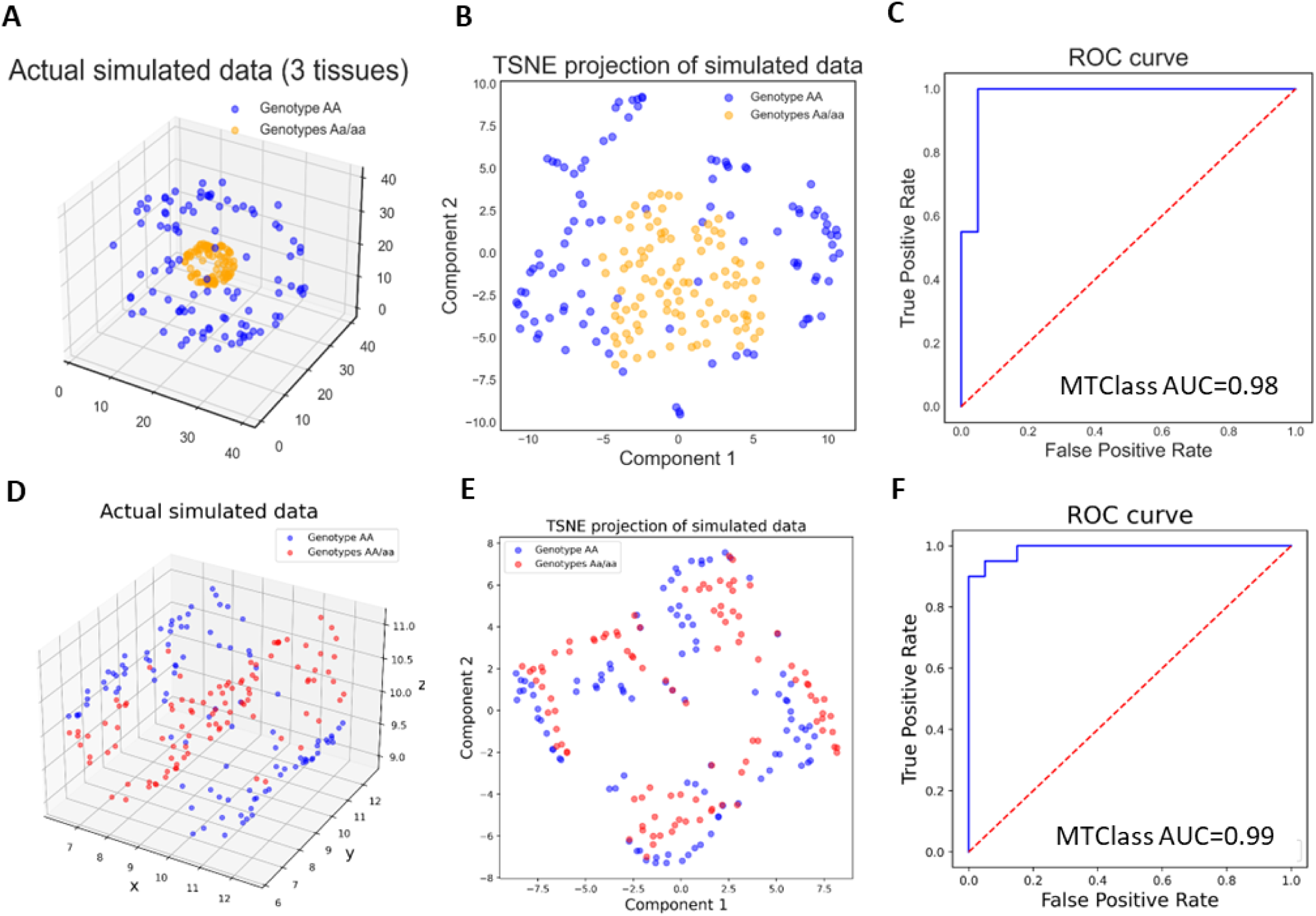
Simulation study using nonlinear genotype-phenotype relationships to demonstrate the utility of MTClass. **A-C** and **D-F** represent two simulated datasets, each with 100 samples for 2 classes, where the genotypes were not linearly separable in two-dimensional space. MTClass was still able to achieve high AUROC in both cases. **A.** Visualization of spherical simulated dataset with differences in variance between two genotypes. **B.** t-SNE dimensionality reduction of simulated data shows nonlinear separability between the two genotypes. **C.** MTClass achieves high AUROC even with nonlinear separability between the two classes. **D.** Visualization of sinusoidal simulated dataset. **E.** t-SNE projection of second simulated dataset. **F.** MTClass also achieves very high AUROC with this dataset.

**Supplementary Figure 2.**
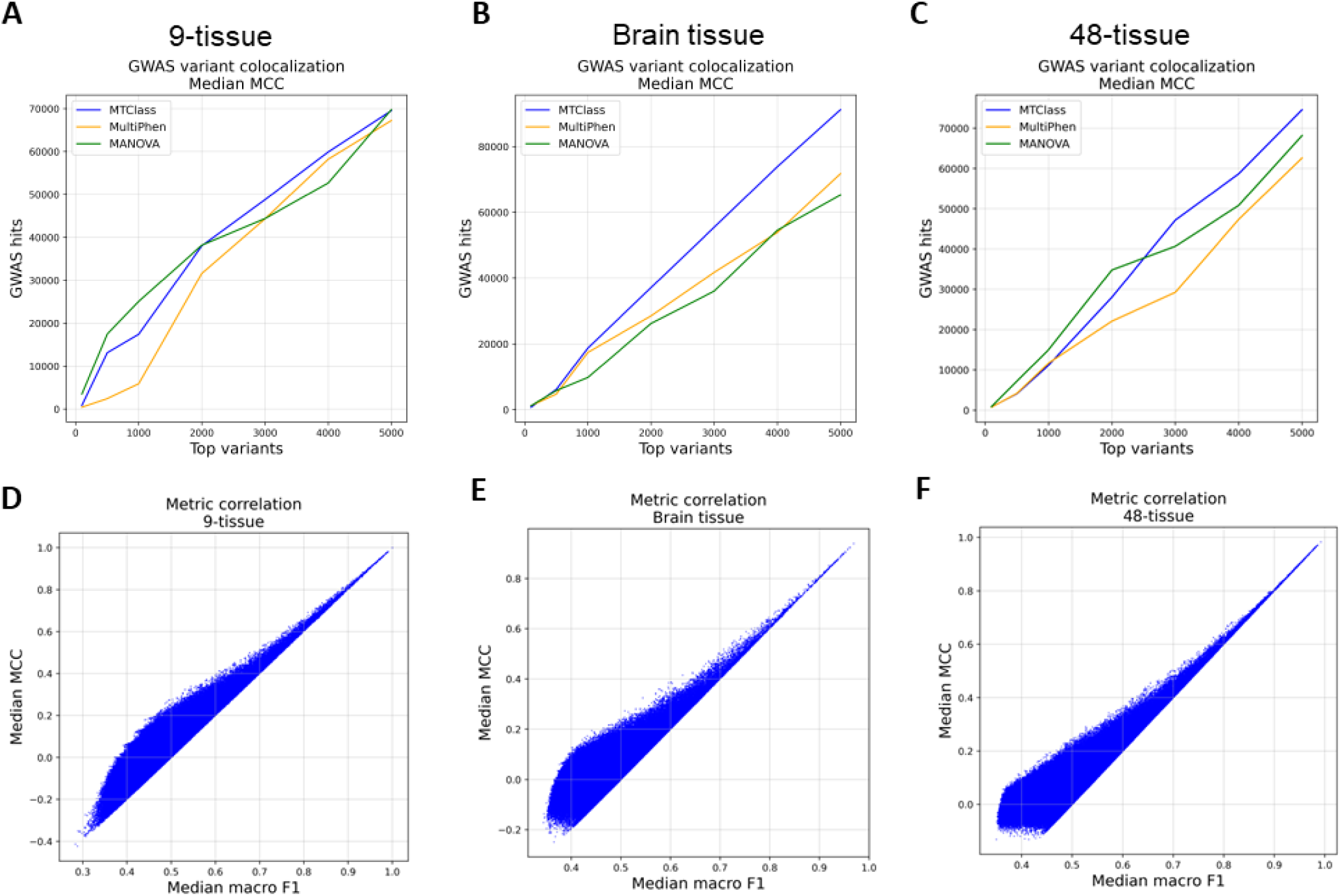
**A-C.** GWAS variant colocalization analysis using the median Matthews correlation coefficient (MCC) across 3 iterations produces similar results as using median macro F1 score in **A.** the 9-tissue case, **B.** the brain tissue case, and **C.** the 48-tissue case. **D-F.** Correlation between the median macro F1 score and median MCC score in **D.** the 9-tissue case, **E.** the brain tissue case, and **F.** the 48-tissue case. In all cases, the top variants showed particularly high correlation between these two metrics.

**Supplementary Figure 3.**
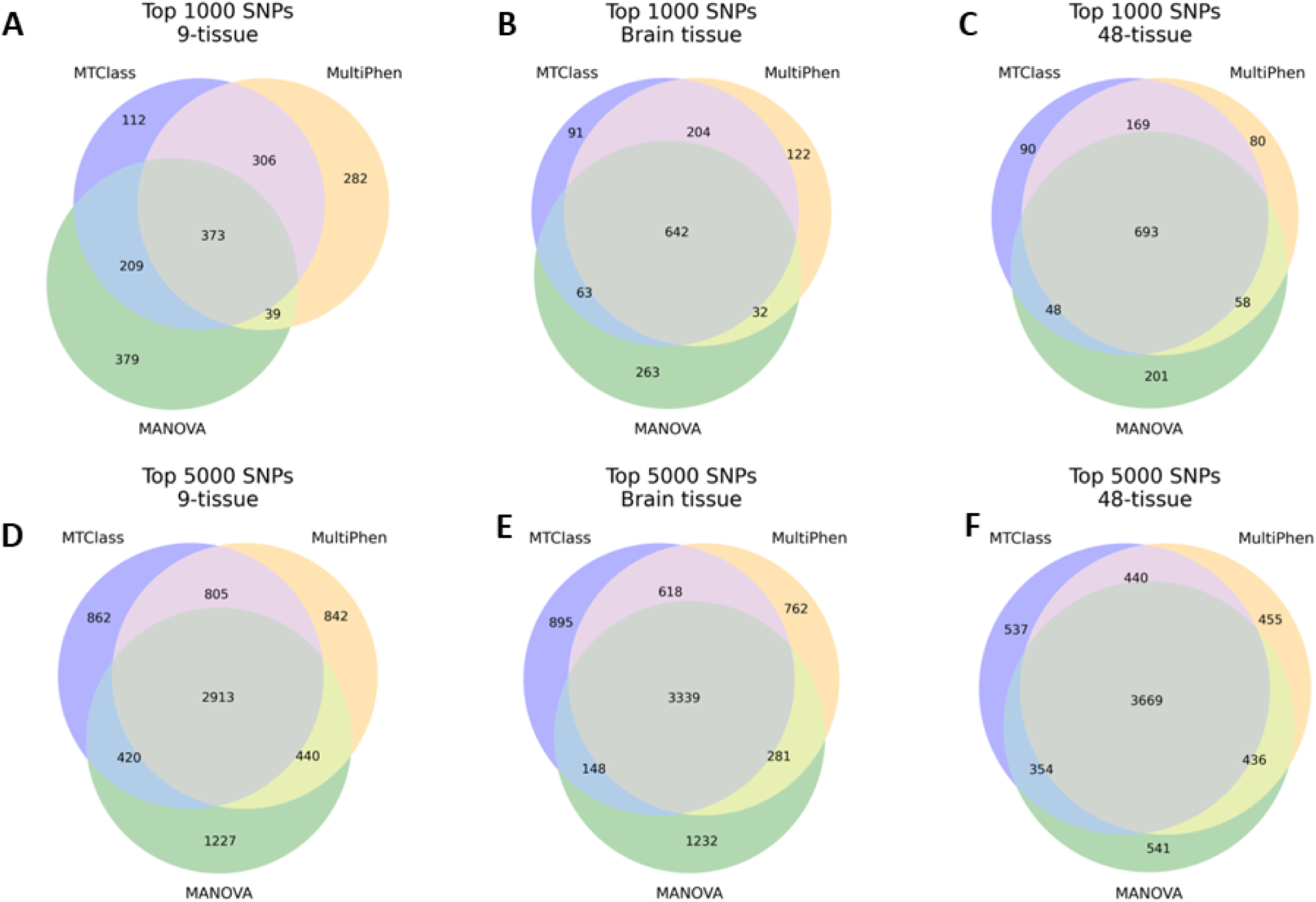
**A-C.** Overlap among the top 1000 variants for the **A.** 9-tissue case, **B.** brain tissue case, and **C.** 48-tissue case. **D-F.** Overlap among the top 5,000 SNPs for the **D.** 9-tissue case, **E.** brain tissue case, and **F.** 48-tissue case. Overall, there is a significant amount of overlap in the top variants detected among the three methods. However, each method still identifies a substantial number of unique variants.

**Supplementary Figure 4.**
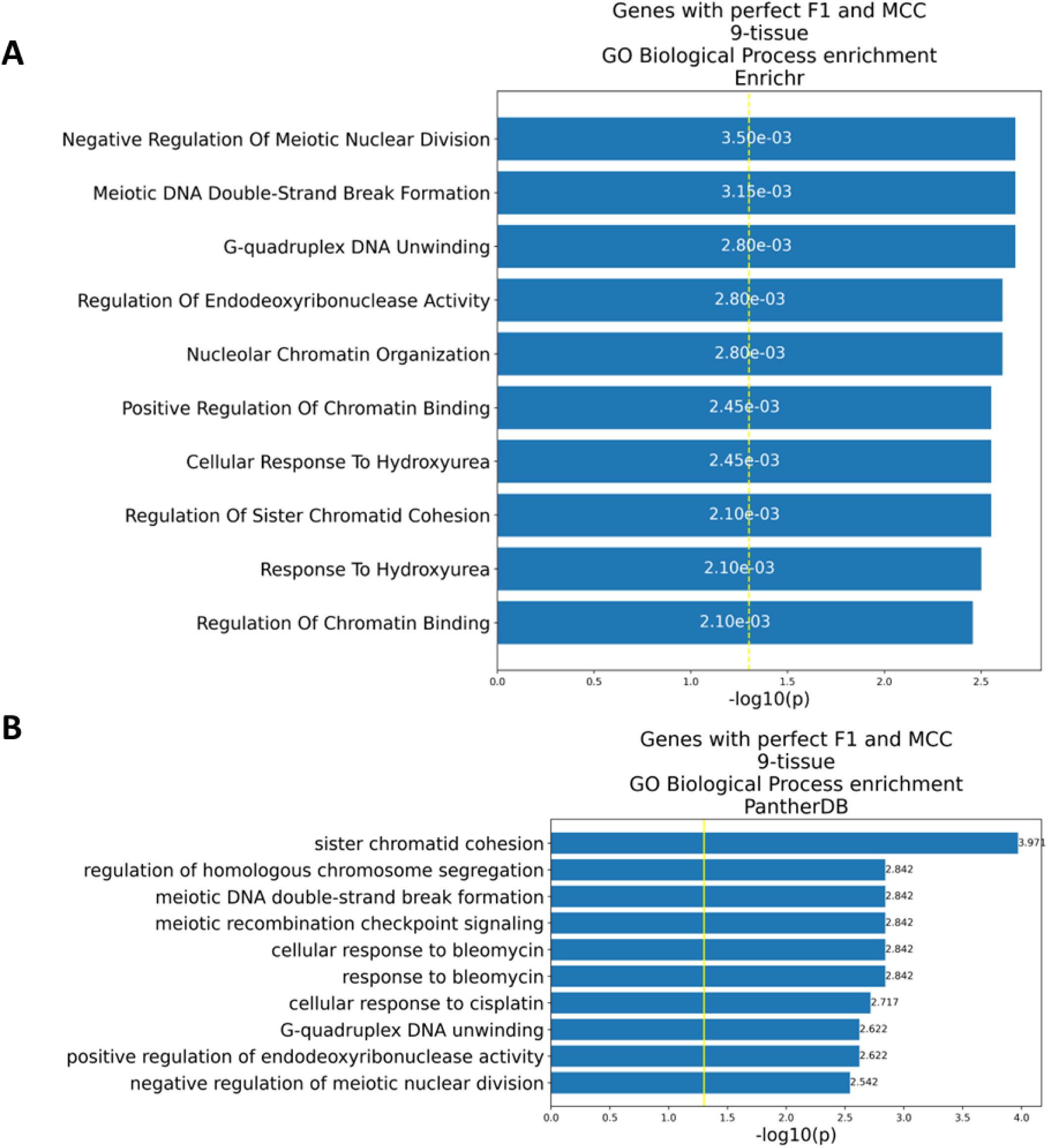
Gene set enrichment analysis of the 8 genes in the 9-tissue case that contained variants with perfect classification performance. These genes are enriched for nuclear and cell division processes. **A.** Enrichment analysis of the 8 genes in the 9-tissue case using Enrichr. **B.** Enrichment analysis of the same genes using PantherDB.

**Supplementary Figure 5.**
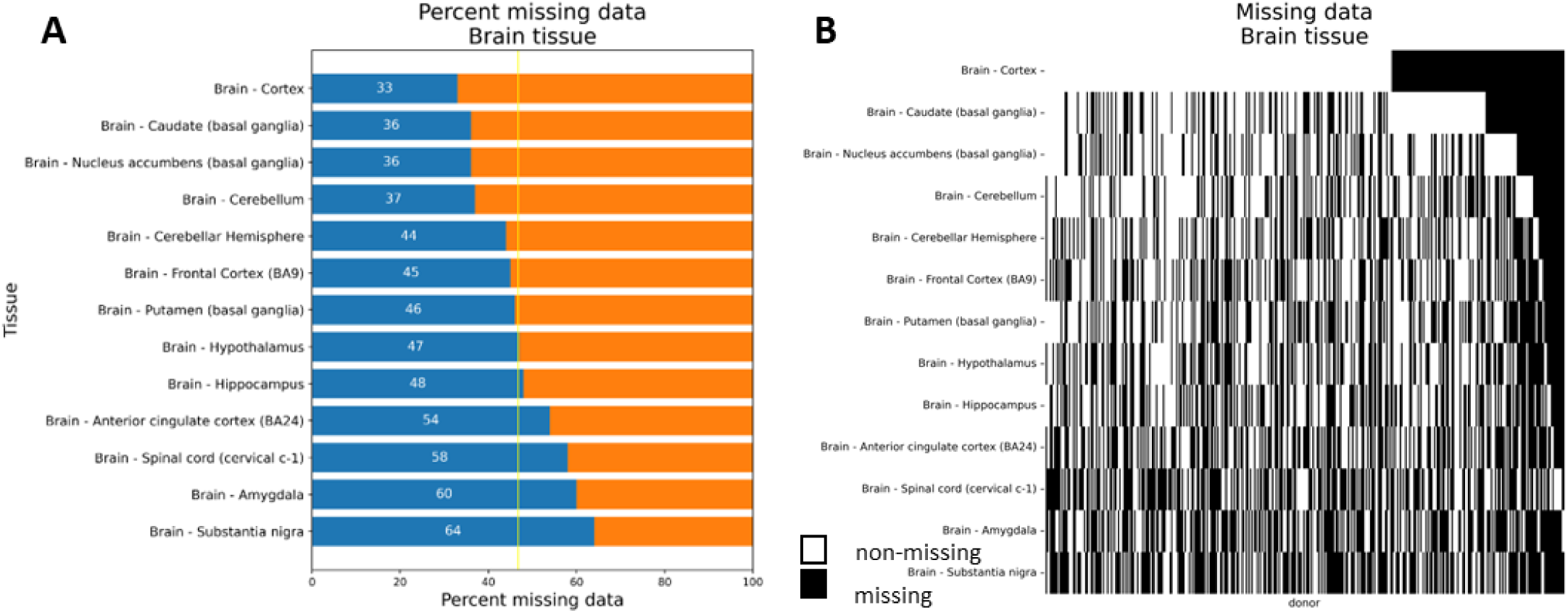
Visualization of missing data in the multi-tissue, brain tissue case. The brain tissue case was missing approximately 47% of the data on average. **A.** Percent missing data by tissue. **B.** Heatmap of missing donor-tissue combinations.

**Supplementary Figure 6.**
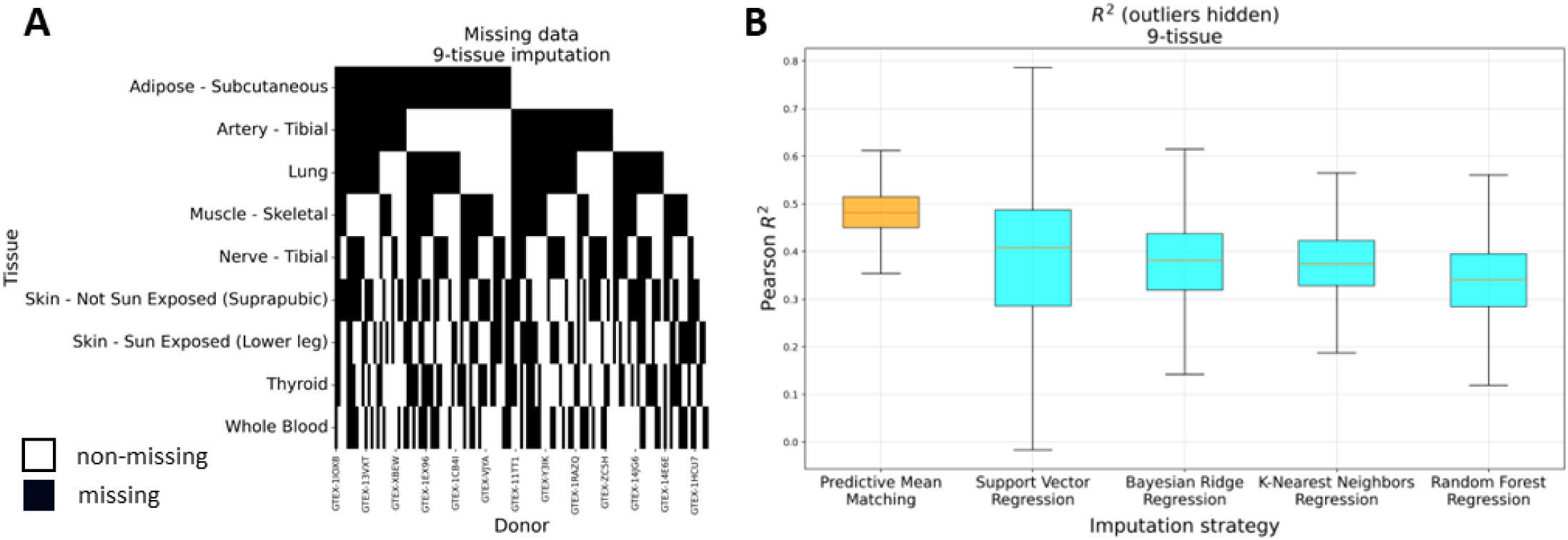
Missing data in the 9-tissue imputation study. 50% of the donor-tissue combinations were randomly removed from the 9-tissue case, where the ground truth is known, to test various imputation strategies. **A.** Heatmap of randomly removed donor-tissue combinations. **B.** Boxplot of Pearson correlation coefficient between true and imputed gene expression measures by imputation strategy. Predictive mean matching had the highest median *R*^2^.

**Supplementary Figure 7.**
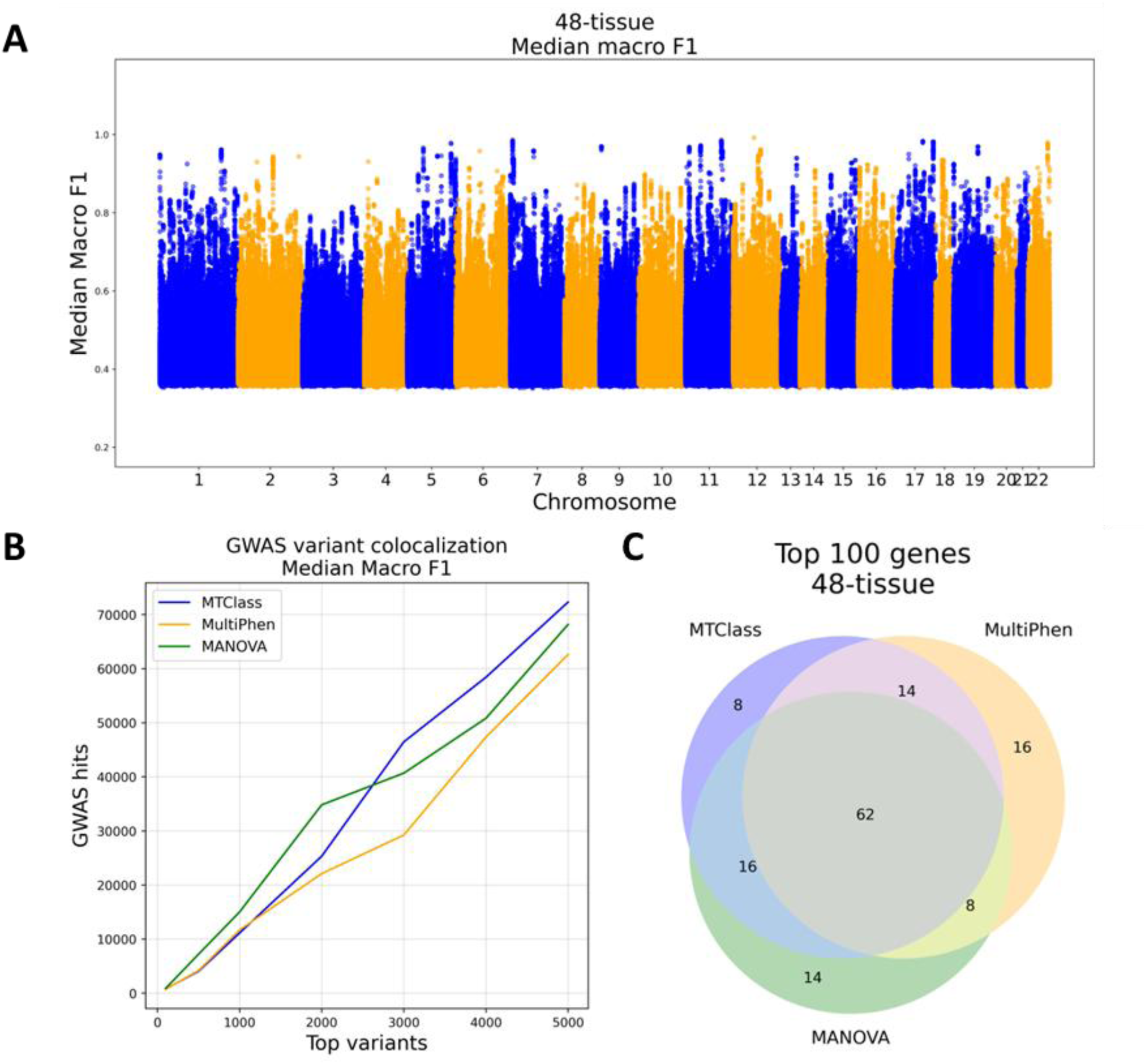
Results from the 48-tissue case (multi-tissue study). **A**. Manhattan plot of variants according to their median macro F1 score. **B.** GWAS variant colocalization analysis of top variants from each method. **C.** Overlap of top 100 genes detected by each method.

**Supplementary Figure 8.**
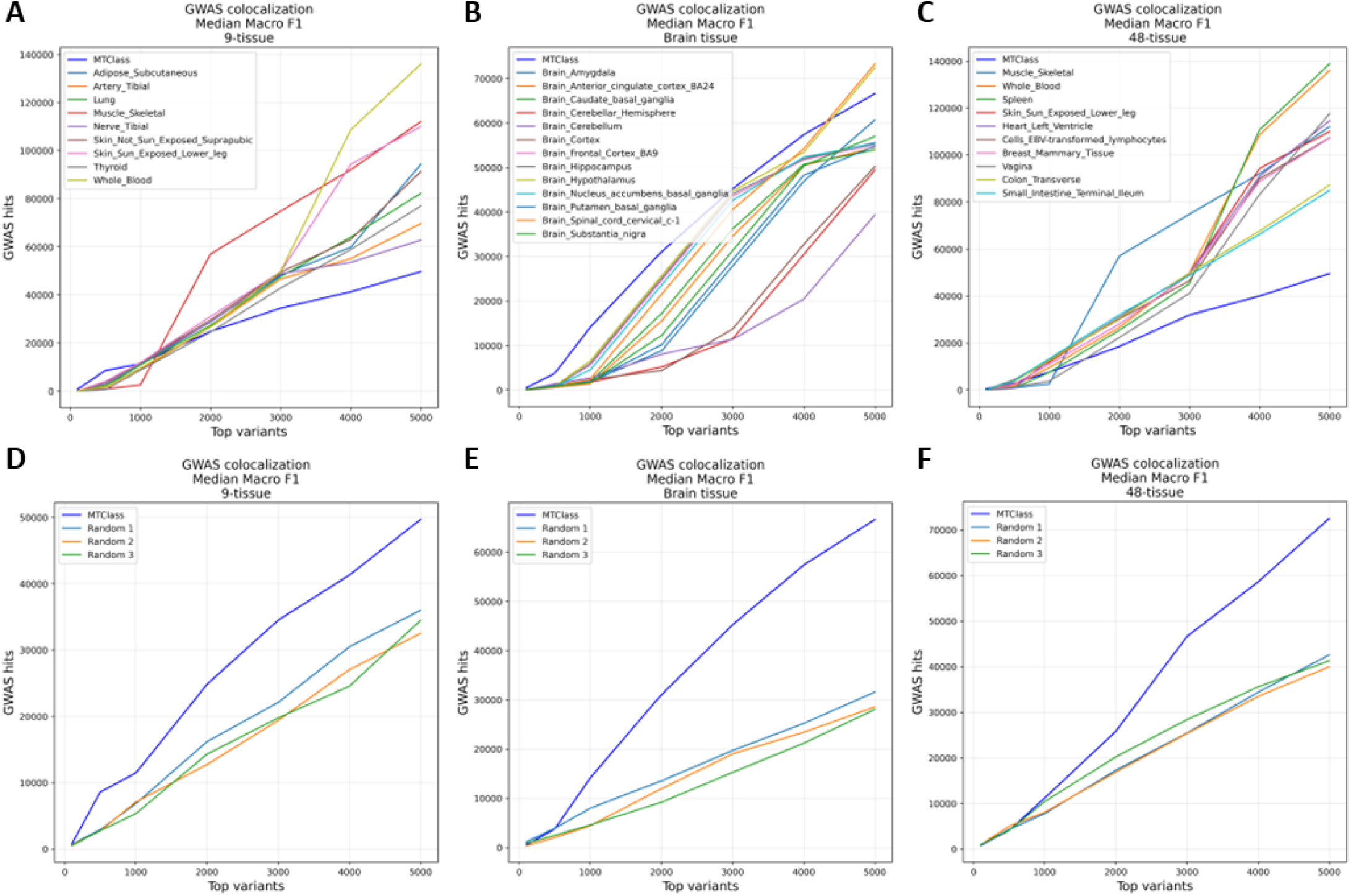
**A-C.** GWAS variant colocalization analysis compared to the top single-tissue eQTLs in the **A.** 9-tissue, **B.** brain tissue, and **C.** 48-tissue cases. In the 48-tissue case, only the top 10 tissues with the highest average GWAS hits were plotted for ease of visualization. **D-F.** GWAS variant colocalization analysis compared to three random subsets of eQTLs taken from all tested eQTLs in the **D.** 9-tissue, **E.** brain tissue, and **F.** 48-tissue cases. MTClass always achieves more GWAS colocalization than randomly selected subsets of results, but not compared to the single-tissue eQTLs in the 9-tissue and 48-tissue cases.

**Supplementary Figure 9.**
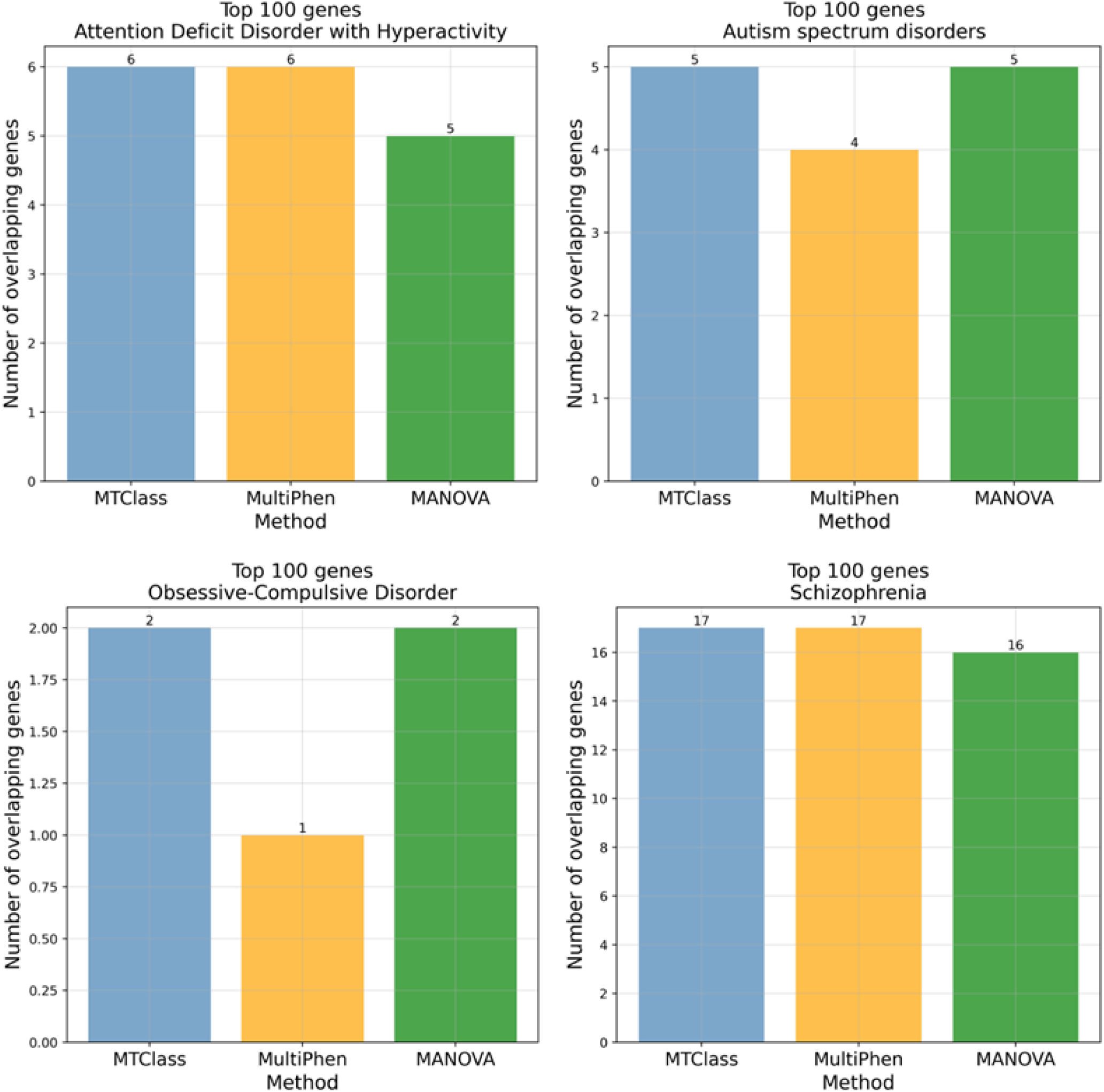
Disease-associated gene overlap analysis in the PsychENCODE isoQTL study with other prefrontal cortex-related traits such as attention deficit-hyperactivity disorder (ADHD), autism spectrum disorder (ASD), obsessive-compulsive disorder (OCD), and schizophrenia. Although MTClass does not always singly detect the greatest number of disease-associated genes, our method always ties with the leading method in the analysis.

**Supplementary Figure 10.**
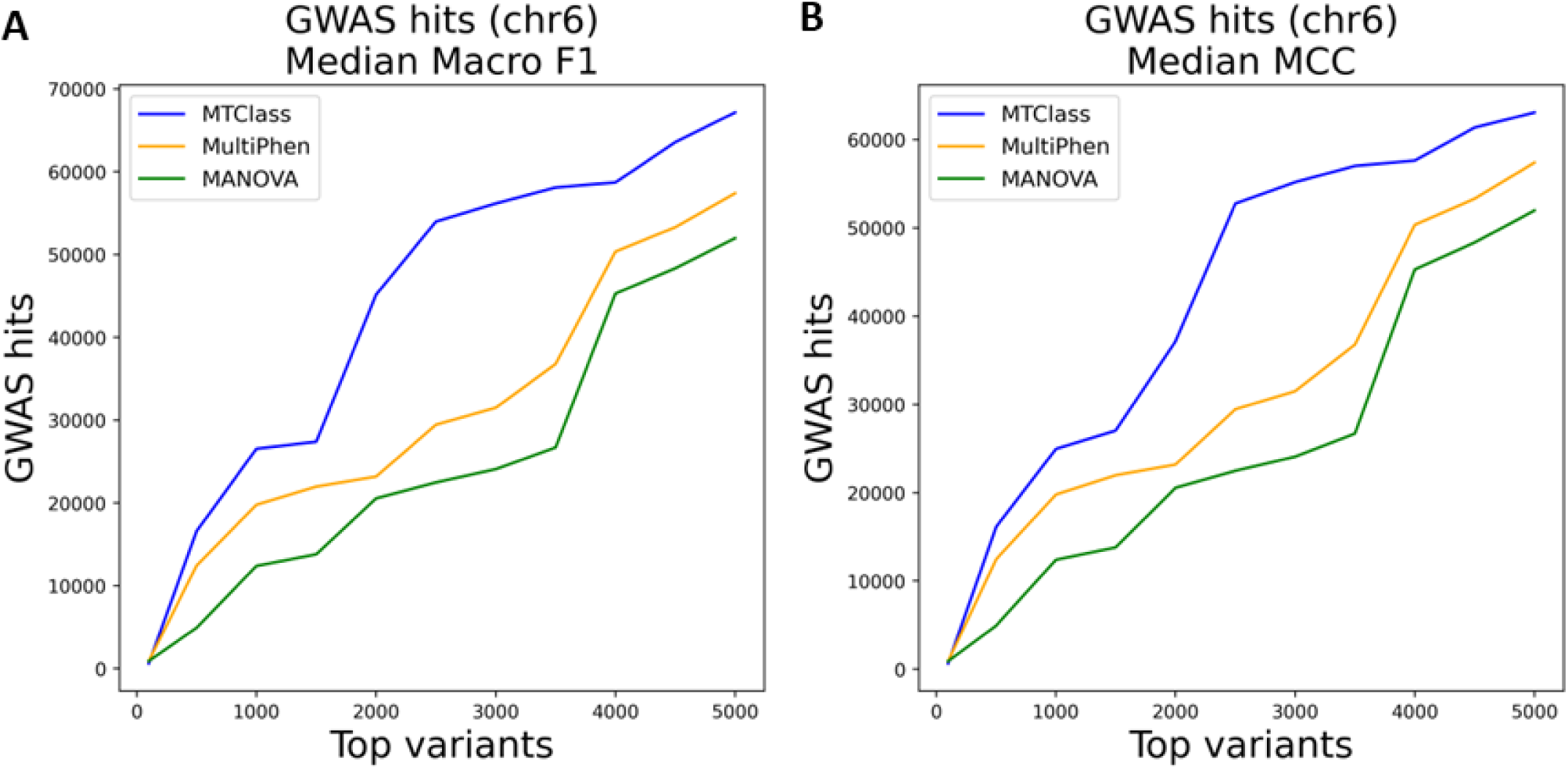
GWAS variant colocalization analysis on the OneK1K scRNA-seq cohort shows that hits on chr6 contribute to the majority of the total GWAS hits among the top variants, after sorting by both **A.** median macro F1 and **B.** median MCC. Due to the abundance of HLA genes on chr6, we concluded that immune-related features were of great importance, especially to MTClass.

**Supplementary Figure 11.**
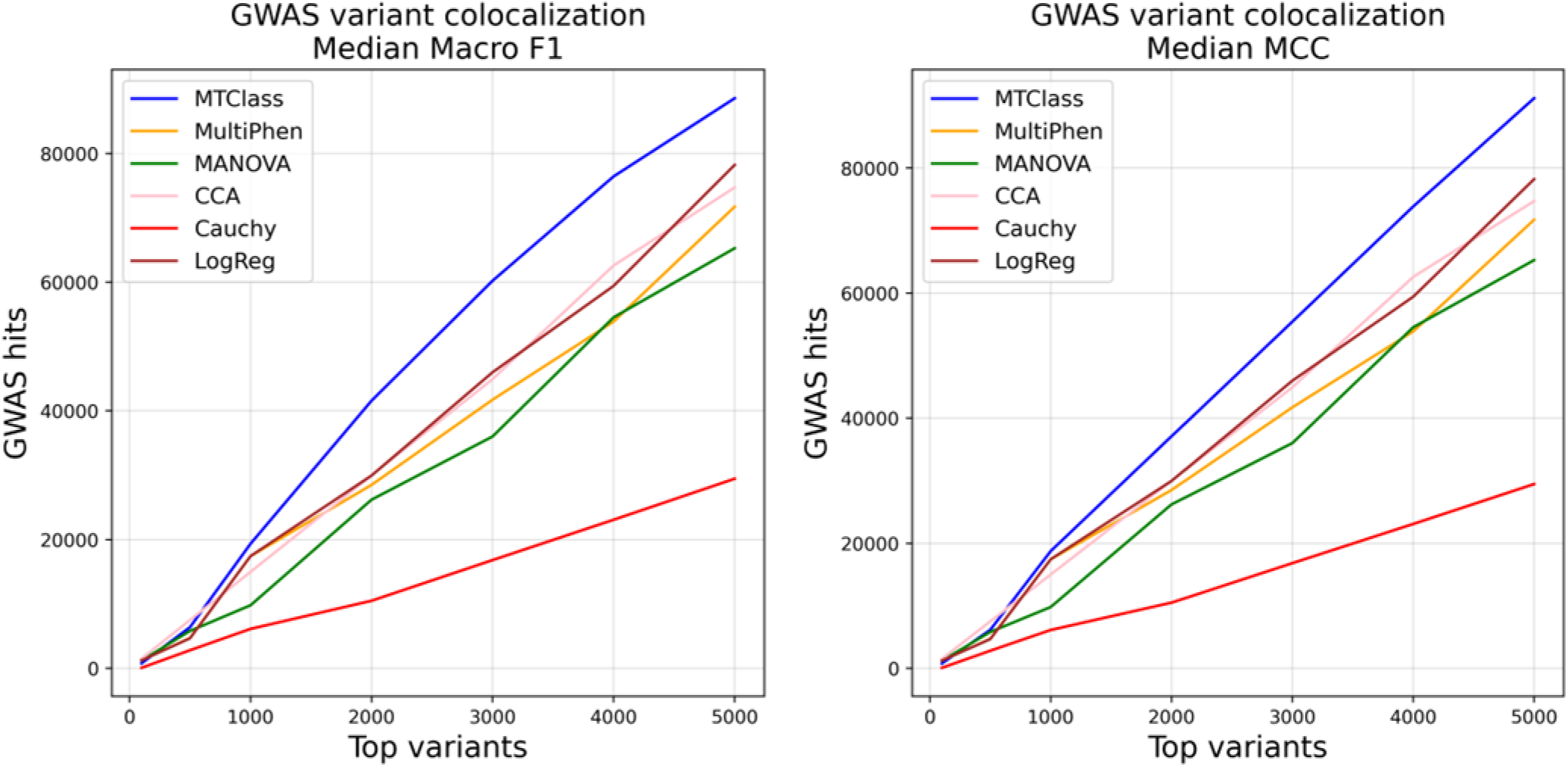
Extended performance comparison with more multivariate association methods in the brain tissue case shows that MTClass is superior at detecting eQTLs of high functional importance. In addition to MultiPhen and MANOVA, we compared MTClass to canonical correlation analysis (CCA), reverse logistic regression, and the Cauchy combination test.

